# Cancer cells communicate with macrophages to prevent T cell activation during development of cell cycle therapy resistance

**DOI:** 10.1101/2022.09.14.507931

**Authors:** Jason I. Griffiths, Patrick A. Cosgrove, Eric Medina Castaneda, Aritro Nath, Jinfeng Chen, Frederick R. Adler, Jeffrey T. Chang, Qamar J. Khan, Andrea H. Bild

**Affiliations:** Department of Medical Oncology & Therapeutics Research, City of Hope National Medical Center, 1500 East Duarte Road, Duarte, CA, 91010, USA; Department of Mathematics, University of Utah 155 South 1400 East, Salt Lake City, UT, 84112, USA; Division of Medical Oncology, Department of Internal Medicine, The University of Kansas Medical Center, Kansas City, KS, 66160, USA; State Key Laboratory of Integrated Management of Pest Insects and Rodents, Institute of Zoology, Chinese Academy of Sciences, Beijing, China; School of Biological Sciences, University of Utah 257 South 1400 East, Salt Lake City, UT, 84112, USA; Department of Integrative Biology and Pharmacology, School of Medicine, School of Biomedical Informatics, UT Health Sciences Center at Houston, Houston, TX, 77030, USA

## Abstract

Cancer cells evolve, acquire resistance to therapy and change their environment. One resistance mechanism involves altering communication with non-malignant cells in the tumor microenvironment (TME). By corrupting the signals for growth and survival, evolving cancer cells can engineer a pro-tumor TME. However, the specific interactions between malignant and non-malignant cells that predispose drug resistance and their changes during treatment remain widely unknown. Here we examine the composition, communication, and phenotypic diversity of tumor-associated cell populations in serial biopsies from early-stage ER+ breast cancers. These patients received either endocrine therapy (letrozole) alone or in combination with a CDK4/6 cell cycle inhibitor ribociclib, and were analyzed using single-cell RNA sequencing (scRNAseq). Our analyses reveal cancer cells from ribociclib resistant tumors stimulate macrophage differentiation towards an immune-suppressive phenotype through upregulation of a broad diversity of cytokines and growth factors. This shift in phenotype leads to reduced macrophage cell IL-15/-18 crosstalk with CD8+ T-cell via IL-2/15RA/18R receptors, resulting in diminished T-cell activation and recruitment. Thus, cancer communication promoting an immune-cold TME predispose tumors to develop CDK4/6 inhibitor resistance, and that the beneficial effects of cell cycle inhibitors through blocking cancer cell proliferation must be balanced against their known inhibitory effect on immune cell division and activation. An optimal treatment strategy will require coupling the prevention of cancer division with activation of an effective cytotoxic T-cell response.

## Introduction

In healthy tissues, interactions among epithelial, stromal and immune cells tightly regulate cell phenotypes, proliferation and tissue composition ^**1**^. These cellular communication networks are disrupted in tumors, with a strengthening of growth-promoting signals and weakening of growth-inhibitory controls ^2–4^. The milieu of communications between cancer and non-cancer cell types impact the onset of malignancy and engineer the tumor microenvironment (TME) to allow disease progression and establishment of a metastatic niche ^5^.

The phenotype, communication and composition of non-cancer cells within the tumor influence treatment resistance, in addition to the genetic heterogeneity and evolution of cancer cells. Diverse non-cancer cell types can modulate growth and survival signals in the TME, potentially contributing to resistance and progression. For example, tumor-associated macrophages can differentiate into an immune-suppressive M2 phenotype instead of an immune activating M1 state, switching signals in the TME from an anti-cancer to a pro-cancer state ^6^. Additionally, fibroblasts can promote extracellular matrix deposition to support cancer cell growth, while endothelial cells can support angiogenesis to supply oxygen and nutrients to a growing tumor ^7,8^. Corrupt cancer cell communications with these non-cancer cells of the TME allow exploitation of their regulatory functions to engineer a pro-tumor TME or avoid immune surveillance ^9^.

Our recent studies show that early-stage estrogen receptor positive (ER+) breast cancer cells upregulate growth factor receptors to amplify alternatives to estrogen growth signaling after treatment with endocrine and cell cycle therapy to bypass cell cycle arrest and promote resistance^10^. Targeting such aberrant communications provides therapeutic opportunities to block tumor-promoting TME interactions and control cancer proliferation ^5,11^. However, it remains unknown how heterogeneity in tumor communication, composition and cell phenotypes prior to and during treatment regulate tumor response to specific treatments. Additionally, research is needed to determine if cancer cells can modify non-cancer regulatory communications to produce a supportive TME and treatment resistance.

Communications are often mediated by ligand-receptor (L-R) interactions. Ligand signals produced by diverse cell types accumulate in the TME and bind to receptors on receiving cells. Signal transduction to the nucleus controls gene expression, culminating in changes in cell phenotype and function. Insights into how cellular phenotype influences communication between individual pairs of cells can be obtained by deciphering cell–cell interactions (CCI’s) from transcriptomic data ^12^. Individual level CCI’s are then inferred from ligand and receptor gene expression of the sending and receiving cell and tested using permutation ^13–16^ or graph-based approaches ^17–19^. However, the ability of cancer and non-cancer cell types to amass corrupting signals in the TME depends on the cellular abundance and composition of each phenotype, with rare cell types contributing sparse signals across the TME, even if individual cells are active communicators.

To understand how phenotypically diverse populations of cancer and non-cancer cells in a tumor communicate through contribution and receipt of signals, we developed TWISTER (**T**umor-**W**ide **I**ntegration of **S**ignaling **T**o **E**ach **R**eceiver). TWISTER extends the individual level CCI concept to measure population-level signaling from across all single cells profiled in a tumor (i.e. tumor-wide). This uncovers networks of communication between the phenotypically diverse populations of cancer and non-cancer cell types constituting the tumor, accounting for both composition and phenotype. TWISTER measures tumor-wide communications received by each cell from diverse cancer and non-cancer cell populations using single-cell RNA sequence (scRNAseq) transcriptomic profiles and detailed annotations of the tumor’s cell type composition. This tumor-wide perspective of communication is essential to study the cancer ecosystem as a whole. Different cell subtypes can have conflicting roles in TME engineering and the abundance and strength of signaling of each cell population influences the tumor progression. One example is the relative abundance of the dichotomous tumor-promoting and suppressing M1/M2 macrophage populations respectively ^20^.

Here we measure communications between non-cancer and cancer cell populations in early-stage estrogen receptor positive (ER+) breast cancer tumors. Using serially collected biopsies from 63 patients taken prior to, during and after treatment, our studies determine how communication differs between tumors resistant and sensitive to endocrine and cell cycle inhibitor therapies. The diversity of cell populations and their communications in sensitive and resistant tumor microenvironments (TME) is measured to reveal ecosystem-wide divergence between their composition and communication, and shows how cancer signaling to macrophages impairs T cell recruitment and activation and drives resistance to cell cycle inhibitor treatment.

## Results

### Patient treatment, sample collection and tumor response

We studied the tumor-wide communication among cells in tumors of post-menopausal women with node positive or >2 cm ER and or PR+, HER2 negative breast cancer enrolled on the FELINE clinical trial ^10,21,22^(clinicaltrials.gov # NCT02712723). This trial evaluated the efficacy of combining CDK inhibition with endocrine therapy in the neoadjuvant setting. Tumor-wide communication was determined in patients randomized to receive either combined CDK inhibition and endocrine therapy (combination ribociclib= ribociclib + letrozole) (n=80 patients) or endocrine therapy alone (letrozole alone= letrozole + placebo) (n=40 patients). Patients were treated for six months and biopsies were collected at baseline (day 0), following treatment initiation (day 14), and end of treatment (surgery around day 180). Each patient’s tumor was identified as growing or shrinking during therapy, using multi-model tumor growth measurements over time (detailed in ^10^). Growing tumors resistant to therapy had a higher proportion of tumor remaining after therapy (>2/3 initial size) (t-statistic=4.45, p<0.001) and classifications were well aligned with clinical assessments.

### Discovery and validation cohort sequencing

The 120 patients were divided into two equally sized cohorts; a hypothesis generating discovery cohort and a validation cohort. Two-thirds of the patients in each cohort received the combination ribociclib treatment, while the remainder received letrozole alone. Single-cell RNA sequencing (scRNAseq) was performed on each serially collected sample of the tumors (detailed in ^10^). In the discovery cohort, 35 patients provided high-quality biopsy samples yielding serial time-point scRNAseq (10X) data for analysis of cell type, phenotype, communication and composition (**Figure 1A**). Of these patients, 23 received combination ribociclib (13 resistant and 10 sensitive tumors) and 12 received letrozole alone (5 resistant and 7 sensitive tumors). The validation cohort was sampled and processed following the same procedures and we additionally rescued some lower quality cells so as to retain a greater number of non-cancer cell types (especially immune cells) across samples. From the validation cohort biopsies, high-quality serial scRNAseq data were obtained for 27 patients, of which 17 received combination ribociclib (5 resistant and 11 sensitive tumors) and 11 received letrozole alone (7 resistant and 4 sensitive tumors).

**Figure 1.**
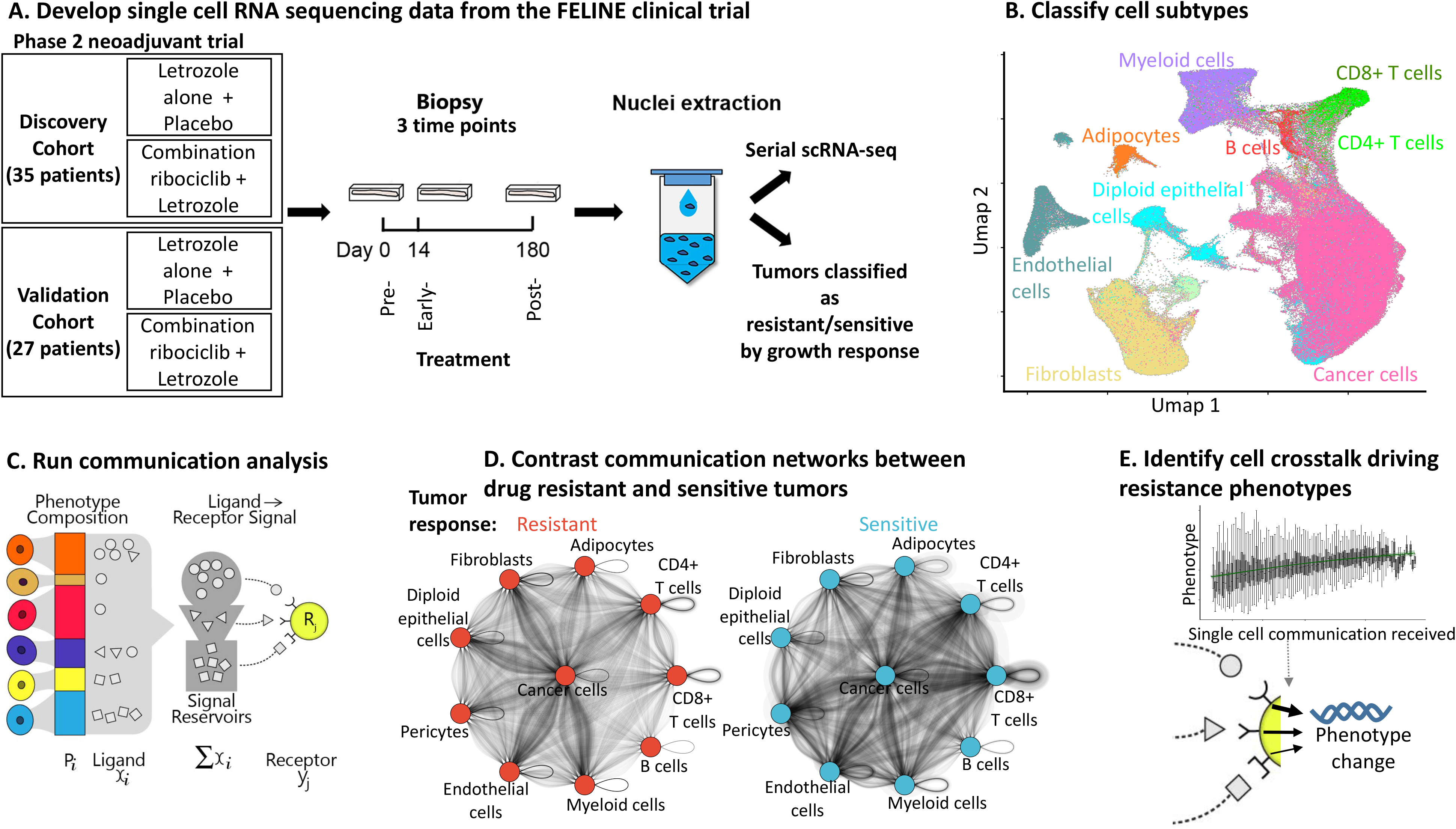
Composition and communication of phenotypically diverse cancer and non-cancer cells within the tumor microenvironments of early-stage ER+ breast cancer patients resistant or sensitive to cell cycle or endocrine therapy. A) Schematic diagram of clinical trial design and single- cell RNA-seq workflow. Serial single cell data was generated for 63 patients using 10X genomics. A total of 424,581 high quality cells were transcriptionally profiled. B) Distinction of cancer and non-cancer cell types shown by UMAP dimension reduction plot of single cell gene expression profiles (points= single cells, color=cell type). C) Overview of the TWISTER algorithm used to measure networks of communication between the phenotypically diverse populations of cancer and non-cancer cell types constituting the tumor, accounting for both composition and phenotype. Phenotypically diverse subpopulations of each cell type were resolved using UMAP and relative abundance calculated in each tumor sample (proportion subpopulations i= Pi). For each subpopulation (cell color), ligand expression (shapes) quantifies the per cell signal production and the total across the subpopulation of cell indicates the contribution to signals in the TME. For each signal receiving cell, the corresponding receptor expression quantifies the ability of each single cell to receive signals from the TME. D) Networks of communication between cancer and non-cancer cell types via diverse ligand-receptor communication pathways in resistant (red) and sensitive tumors (blue). Nodes (circles) indicate cell type and edges (weighted arrows) indicate the strength of communication between a pair of cells (dark lines= strong communication). Arrows are directed, with curves going from the sender to the receiver in clockwise direction. E) Single cell communication received was associated with cellular differentiation and activation of a resistance related phenotype in cancer or non-cancer cells.

### Cell type annotation and verification

We obtained high quality transcriptional profiles for 424,581 single cells (41% discovery cohort, 59% validation cohort) with stringent quality controls ensuring high-coverage, low mitochondrial content, and high-confidence of doublet removal. For each cohort, broad cell types were annotated using singleR ^23^, and cancer cells were identified by their frequent and pronounced copy numbers amplification using inferCNV ^24^. Cell type annotations were verified by cell type specific marker gene expression and Umap/TSNE analyses (**Figure 1B**) ^10,25^. Granular immune subtype annotations were obtained using our recently published ImmClassifier machine learning method, which has been validated against flow cytometry assays ^26^. Cell subtype annotations for the discovery and validation cohorts were confirmed to be consistent by training a random forest machine learning classifier to predict cell types using the discovery cohort data and then applying it to predict cell annotation in the validation cohort (see methods). We obtained a high agreement of >75% between annotation approaches (Figure S1).

### Communication between phenotypically diverse populations

To understand how the TME composition and communication can contribute to resistance, we applied TWISTER to measure tumor-wide signaling from diverse non-cancer cell sub-populations and heterogeneous cancer lineages to each receiving cell (**Figure 1C**). existing cell-cell interaction approaches reveal how one cell of one type communicates with another. Crucially, our generalization measures the signal each individual cell receives from many phenotypically diverse subpopulations of cells (e.g. signal from all M1 vs M2 differentiated macrophages) that all contribute signals to the TME signal reservoir. This accounts for the abundance and ligand production of each signaling phenotype and the receptor activity of receiving cells. To do this, we dissect broad cell types into phenotypically coherent subpopulations and quantify the total contribution of signaling molecules from each group, to reveal how both phenotypic and compositional changes modify communication feedbacks and impact treatment response (**Figure 1D**). We then relate these inferred communications back to observable changes in cellular phenotypes and abundances that influence treatment response (**Figure 1E**).

### Shrinking tumors are enriched with immune cells, while growing tumors are cancer dominated or enriched with stromal and endothelial cells

Prior to examining communication between the cancer and non-cancer populations we compared the relative frequency of each cell type across tumors, allowing identification of archetypical tumor ecosystem compositions observable across early-stage ER+ breast cancers. The compositional similarity of tumor samples was measured using the pairwise Manhattan distance metric. Compositionally similar tumors and cell types with correlated abundances were grouped using hierarchical clustering. Immune cell type abundances were highly correlated with each other; similarly, stromal and endothelial abundances in tumors were correlated (**Figure 2A**). This analysis also indicated that tumors fall into distinct archetypical ecosystem compositions associated with tumor response to treatment.

**Figure 2.**
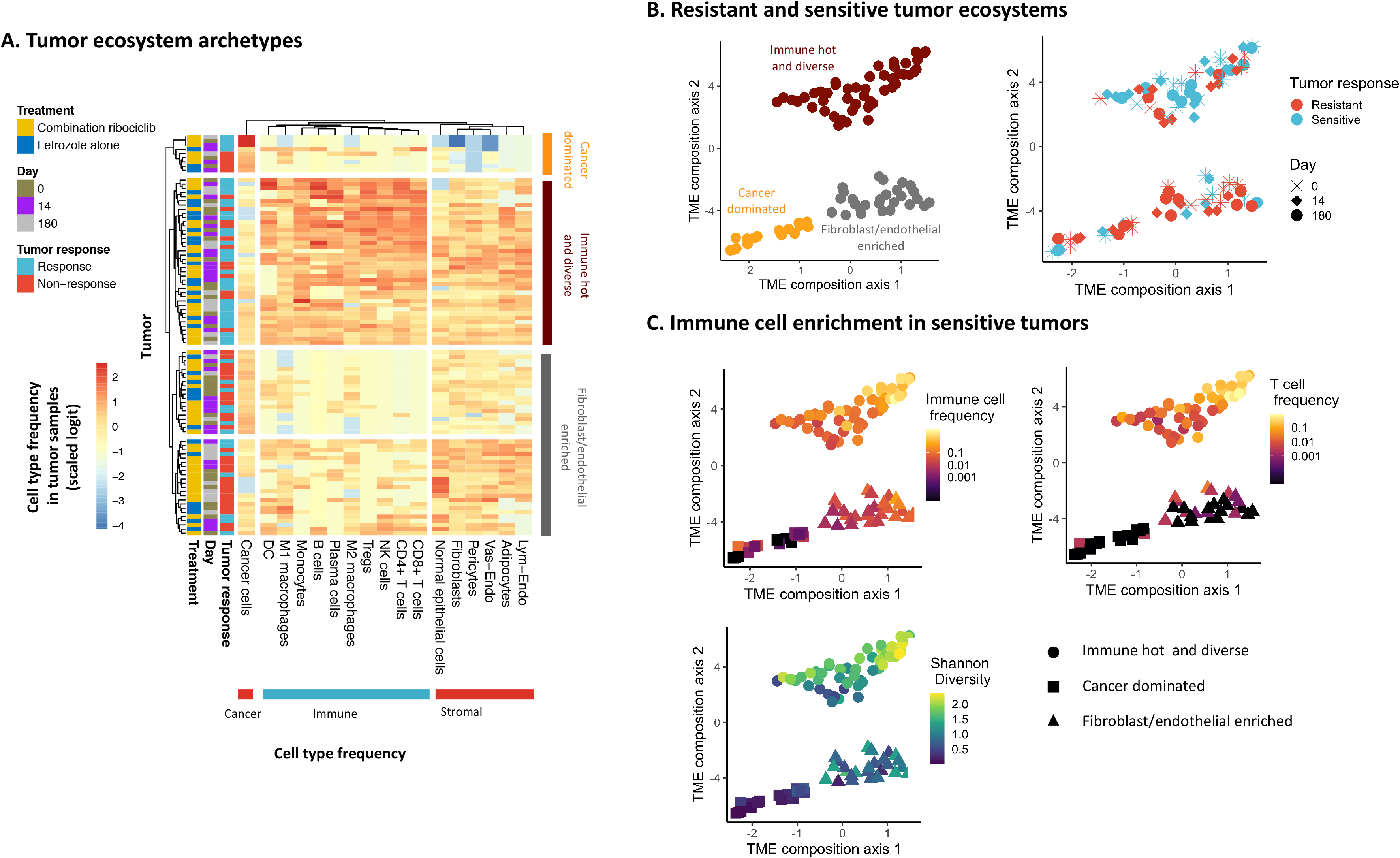
Resistant and sensitive tumors have distinct TME compositions, with sensitive tumors having greater immune cell infiltration. A) Heatmap of the relative abundance of each cell type for each tumor biopsy sample. Tumors are clustered based on pairwise Manhattan distance of compositional similarity. Cell types are similarly clustered to show correlation of pairwise abundance, with immune cell abundances being highly correlated with one another. B) Distinction of three archetypal tumor compositions shown by UMAP dimension reduction using Manhattan distance of compositional similarity (points= tumor samples). Tumor data points close together have high compositional similarity. The distinct archetypal tumor compositions (top left panel colors) were identified using a Gaussian mixture model and then associated with tumor response outcome (top right panel colors). C) Major differences in immune cell type abundance and tumor diversity (Shannon diversity) between archetypal tumor composition states (color=relative abundance, shape= archetype).

We therefore determined the number of tumor compositional archetypes using the UMAP algorithm (Manhattan distance between tumor compositions) (**Figure 2B/C**). Tumors with similar cell compositions clustered close together and distant from tumors with more dissimilar compositions. We identified three archetypal tumor compositions, using a Gaussian Mixture model with a Bayesian information criterion model comparison to determine the appropriate number of TME archetypes. Major compositional differences between archetypal compositions were identified using Dirichlet regression and correlation of TME landscape axes with cell type frequencies. The three distinct tumor archetypal ecosystems (**Figure 2B, left panel**) were: i) a cancer-dominated state, ii) an immune-hot and diverse state, and iii) a fibroblast and endothelial-enriched state. Before treatment, we found resistant and sensitive tumors in all three states, but also a significant association of sensitive tumors with the immune-hot state (association of pre-treatment archetype with tumor response: *χ*^2^=11.285, df=2, p<0.005) (**Figure 2B, right panel**). After treatment, we observed a clear polarization, with 83% of treatment-sensitive tumors in the immune-hot state, and 75% if treatment-resistant tumors in the other two states (Figure S2). This result indicates that resistant breast tumors become increasingly immune-cold and depauperate during treatment. Immune cell loss has been associated with resistance in various cancers ^27^. ER+ breast cancers are often considered uniformly immunologically cold ^28^ even though multiple trials show a subset of ER+ breast cancer patients responding to immunotherapy ^29,30^. T cell abundances were particularly sparse in both non-immune clusters (**Figure 2C, top panels**), highlighting differences between tumor archetypes. In addition, the cancer cell dominant archetype showed a significantly lower Shannon diversity index score (df=89, t=−10.257, p<0.0001) (**Figure 2C, bottom panel**), and was dominated by treatment resistant tumors. In summary, early-stage breast cancer tumors have three main TME compositional archetypes: Immune hot/diverse, fibroblast/endothelial enriched or cancer dominated. Immune hot and diverse tumors, with high T cell and macrophage abundance were more sensitive to cell cycle and endocrine therapy, while immune cold tumors were more resistant.

### Global dysregulation of communication in cell cycle therapy resistant tumors

Communication pathway scores were measured to quantify the communication of ligand signals produced by one cell type population to individual cells of each cell type via a cognate receptor. A total of 1444 ligand-receptor (LR) communication pathways were measured based on known protein-protein interactions (see methods). The overall strength of communication of one cell type with another was determined by averaging across LR communication pathway scores from each sender cell type to the receiver (**Figure 3A/B**).

**Figure 3.**
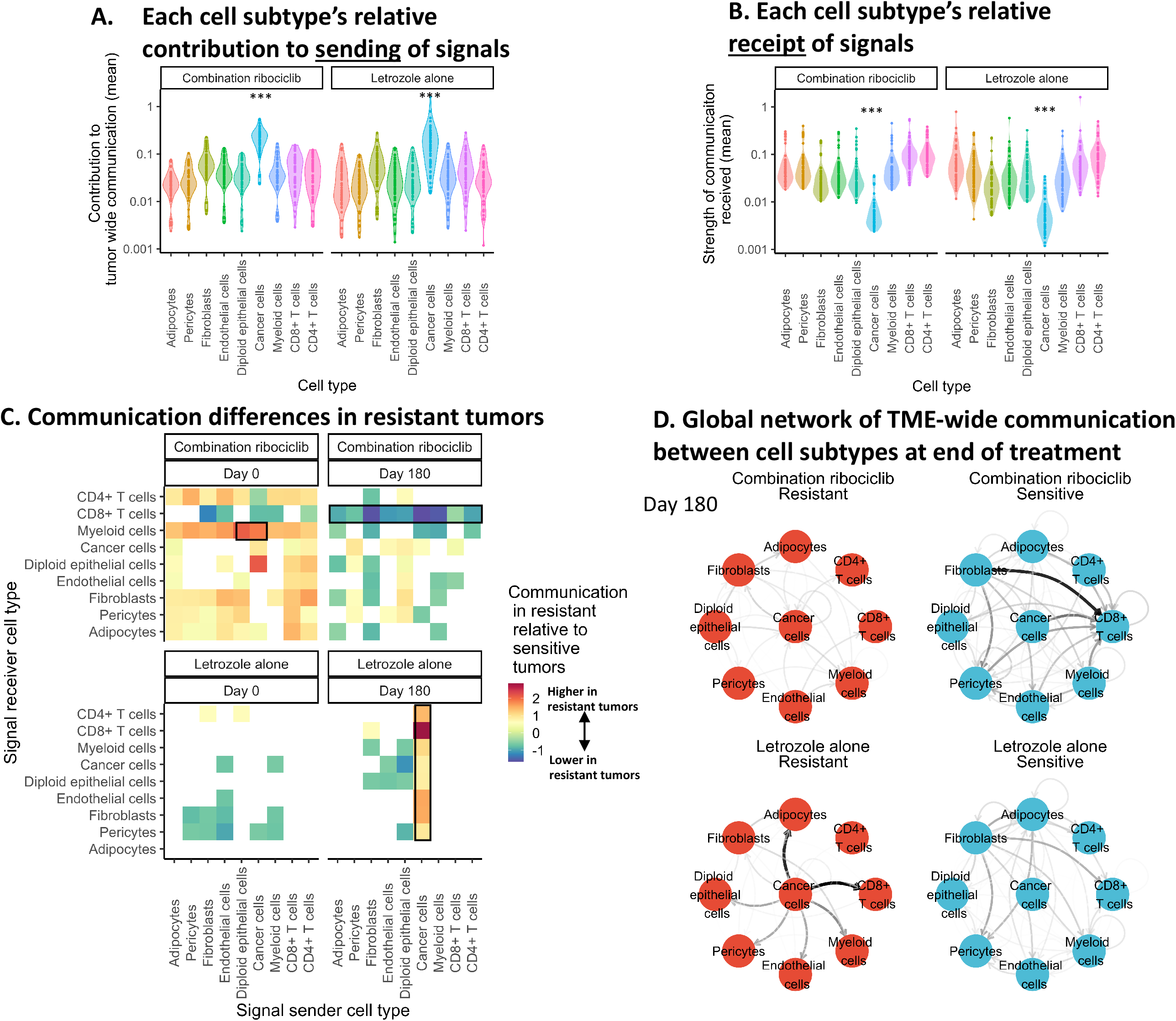
Broad TME wide communication is substantially different across cell types and between resistant and sensitive tumors prior to and after treatment. A) Violin plot showing the contribution of each cell type to sending communications within the TME of tumors given combination ribociclib or letrozole alone. B) Violin plot showing strength of signal received by each cell type. Across tumor samples, cancer cells were consistently the highest contributor to tumor wide signaling but received the least signals. Growth factor signaling via ERBB receptors are an important exception, showing cancer cells do strongly receive a more limited set of signals. C) Differences in the communication from a sending cell population (x axis) to cells of each receiving cell type (y axis) between tumors resistant and sensitive to combination ribociclib or letrozole alone (top vs bottom panel) at the beginning and end of treatment (left vs right panel). Cell types communicating significantly more strongly activated in resistant tumors (red squares) and sensitive tumors (blue squares) were identified using a permutation-based bootstrap randomization. The color intensity indicates the z score of the communication difference in communication difference between resistant and sensitive tumors (white= no significant difference). D) Directed weighted network graphs showing the ecosystem-wide divergence in communication between resistant and sensitive tumors (left vs right panel) following treatment with combination ribociclib or letrozole alone (top vs bottom panel). Cell type represented by nodes and differences in communication were described by the weight of the vertex (arrow) from one cell type to another (arrow width and darkness indicates communication strength).

Across different tumors and treatments, cancer cells contributed more communication signals to the TME than other cell types (**Figure 3A, left panel**) (df=970, t=17.2, p<0.0001). In contrast, cancer cells received the least signal in general from across the TME, receiving substantially fewer communications than non-cancer epithelial, stromal and immune cells (**Figure 3B, right panel**) (df=970, t=−23.52, p<0.0001). However, a small subset of communications, such as growth factor communications (via ERBB family receptors) were most strongly received by cancer cells (df=970, t=10.31, p<0.0001) (Figure S3). These results indicate that cancer cells receive relatively few regulatory signals in the TME while concurrently transmitting broad and strong communications to non-cancer cells in the TME.

We next assessed how communication between cell types differed in therapy resistant tumors before and during treatment. Contrasting communication across many biopsies, rather than within individual biopsies, revealed the TME cell type interactions distinguishing resistant and sensitive tumors and the evolution of communication during treatment. Significant differences in communication between tumors resistant and sensitive to each treatment were identified using a permutation-based bootstrap randomization of the tumor response annotations for each communication pathway (see Methods) (**Figure 3C**). Prior to treatment, the tumors that later developed ribociclib resistance had distinctly different communication networks from those that remained sensitive, showing stronger communication from cancer cells to myeloid cells (**Figure 3C, top left panel**) (df=22, t=3.53, p<0.005) (Figure S4). This strengthening of cancer to myeloid cell communication in ribociclib resistant tumors was verified in the independently profiled validation cohort (z=15.23, p<0.00001). Specific ligand-receptor (LR) communications activated in resistant tumors were identified using log-linear regression with FDR multiple comparisons correction. Most activated LR communication pathways (14/20) bound to myeloid receptors known to promote an immune-suppressive myeloid phenotype (Figure S5) (Table S1:10). These communication pathways were not activated in letrozole resistant tumors (Table S11:20). This result reveals pre-existing communications of cancer cells with myeloid cells that may predispose resistance to cell cycle inhibition but not endocrine therapy.

After 180 days of treatment, cell type communication diverged between resistant and sensitive tumors. In ribociclib resistant tumors, all cell types developed weaker communications with cytotoxic CD8+ T cells (**Figure 3C top right panel**) (3<z<13.5, p<0.0001). In contrast, ribociclib sensitive tumors retained more persistent communications with cytotoxic CD8+ T cells, with stronger signals from fibroblasts and cancer cells (**Figure 3C top right panel**) (z=13.37, p<0.00001). The strong fibroblast-CD8+ T cell interaction reflected increased costimulatory and recruitment integrin communications in ribociclib sensitive tumors at day 180 (e.g. ADAM12-ITGB1:t=4.77, p<0.05)^31,32^. Cancer cells of ribociclib sensitive tumors also provided greater amounts of immune-activating communications to myeloid cells, including stimulation of CCR5/7 receptors ^33^ (df=6, t= 4.73, p<0.01) (Table S2). However, these communication pathways were to expressed at levels too low to be verified in the validation cohort. The tumor-wide decrease in CD8+ T cell communication did not occur during letrozole treatment. In letrozole resistant tumors, cancer cells developed strong communications with stromal and epithelial cells at day 180 (**Figure 3C bottom right panel**) (11<z<31, p<0.0001). The strong cancer-CD8+ T cell communications were regulatory, inhibiting differentiation via TGF*β* and SDC4 signaling (TGF*β*2-TGF*β*R3:t=297.95, p<0.0001; MDK-SDC4: t=225.02, p<0.0001)^34,35^ and promoting exhaustion via CD47 signaling (THBS1-CD47:t=9.03, p<0.01)^36^. Communication between non-cancer cell types became weak. In contrast, letrozole sensitive tumors developed stronger communication between non-cancer cell types, particularly from fibroblasts, with diminished cancer cell signaling.

The communication differences between resistant and sensitive tumors were visualized using directed weighted network graphs for day 180 (**Figure 3D**). Resistant and sensitive tumors showed ecosystem-wide divergence in communication during each treatment (Figure S6), with greater communication between non-cancer cells including T cells in ribociclib sensitive tumors, and stronger communications emerging between cancer and stromal cell types including fibroblasts, endothelial cells and adipocytes in endocrine therapy resistant tumors (**Figure 3D**) (Figure S4). The dynamics of communication reveal the pre-treatment heterogeneity of cancer communication with myeloid cells that predate resistance to cell cycle inhibition but not endocrine therapy, and the breakdown of tumor-regulating immune interactions in ribociclib resistant tumors and an increase of growth-promoting interactions instead in letrozole resistant tumors.

### Corrupt cancer cell communication with myeloid cells prior to treatment stimulates an immune-suppressing macrophage phenotype in ribociclib resistant tumors

As cancer cells of ribociclib resistant tumors communicated immune suppressing signals more strongly to myeloid cells, via a range of LR pathways, we next examined the consequences for myeloid cell phenotype. Myeloid cells are a phenotypically diverse and continuously differentiable population (**Figure 4A, top left panel**) that sense TME conditions and regulate immune responses and wound healing ^7^. Signals from dead cancer cells can promote differentiation to an immune-activating M1 macrophage or dendritic cell phenotype. However, a host of alternative signals, from cancer or non-cancer cells in the TME can promote their differentiation to an immune-suppressive M2 phenotype that supports cancer cell proliferation and survival ^6^. The clear gradient of expression of the CD36 marker of M2 differentiation identifies M2 polarized cells to have high UMAP2 scores (**Figure 4A, top right panel**)^37,38^. Consistent with the detection of immune-suppressive cancer to myeloid communications, we found that macrophages in ribociclib resistant tumors had greater M2 polarization prior to and throughout treatment (df=17.9, t=2.16, p<0.05), while ribociclib sensitive tumors had more M1 macrophages (**Figure 4A, bottom left panel**). Antigen presenting dendritic cells were present in ribociclib sensitive but almost entirely absent from resistant tumors (**Figure 4A, bottom left panel**). The M2 polarization was not present in letrozole resistant tumors (df=10.24, t=1.56, p=0.15) (**Figure 4A, bottom right panel**). One resistant tumor had particularly strongly M1 polarized macrophages. These cells also showed considerable ERBB4 growth factor receptor upregulation which is associated with NRG4 mediated apoptosis of pro-inflammatory macrophage ^39^.

**Figure 4.**
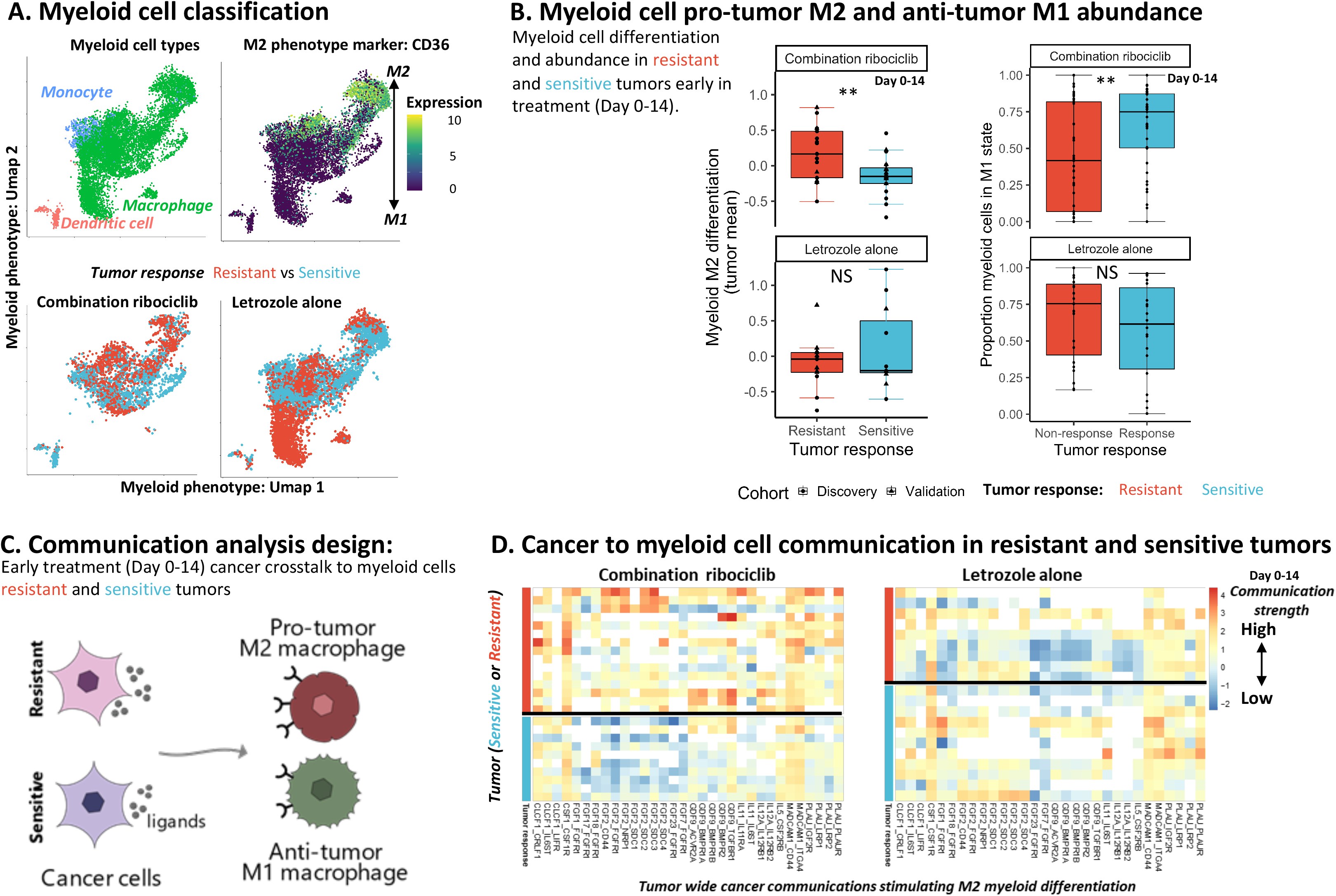
Corrupt cancer cell communications with myeloid cells stimulate pro-tumor M2 macrophage differentiation in resistant tumors. A) Distinction of different myeloid cell subtypes shown by UMAP dimension reduction plot of single cells. Cells with similar transcriptomic profiles are located closer together. Cells with common subtype annotations (obtained using ImmClassifier) clustered tightly in UMAP phenotype space (top left)(color=cell type), showing identifiability of dendritic cell, monocyte and macrophage cell types. The major axis of phenotypic variation within myeloid cells was the gradient of differentiation from an M1 immune-activating phenotype to an M2 pro-tumor phenotype (top right). The gene most differentially expressed across the phenotype space was the M2 phenotype marker gene CD36 (colour gradient= CPM expression). The phenotype of myeloid cells in resistant (red) and sensitive (blue) tumors were compared for patients receiving either combination ribociclib (bottom left) or letrozole alone (bottom right). B) Box and whisker plot (left panel) showing the greater average M2 differentiation of myeloid cells of resistant tumors (red) compared to sensitive tumors (blue) early (Day 0-14) in combination ribociclib treatment (top panel). Black points indicate mean M2 differentiation of specific tumor samples. No significant difference in M2 differentiation was found between letrozole resistant (red) and sensitive (blue) tumors (bottom panel). Box and whisker plot (right panel) showing the lower proportion of immune activating M1 myeloid cells in combination ribociclib resistant tumors. We observed no difference in the proportion of myeloid cell in the M1 state between tumor resistant and sensitive to letrozole. C) Schematic showing the design of the communications pathway analysis used to reveal the activation of M2 differentiation promoting signaling and communication by cancer cells of tumors resistant to combination ribociclib treatment. For each LR communication pathway, the strength of communication sent by the diverse cancer populations of a tumor to myeloid cells was contrasted between treatments resistant and sensitive tumors using log-linear regression. Significant differences in the strength of cell type communication, either before during or after treatment were identified using ANOVA after accounting for multiple comparisons using false discovery rate (FDR) p-value correction. D) Heatmap showing the heterogeneous activity of M2 differentiation communication pathways (columns) across tumors (rows) resistant (red row annotation) and sensitive (blue row annotation) to combination ribociclib (right panel) or letrozole alone (left panel). Heatmap coloration indicates the strength of cancer to myeloid communication within a tumor via a specific M2 differentiation communication pathway (white=no signaling detected). Communication pathways were identified by which cancer cells signal more strongly to macrophages in resistant relative to sensitive tumors or target M2 macrophages (relative to M1 macrophages). Black line separates early treatment (Day 0-14) resistant versus sensitive tumor samples.

To determine whether the pre-treatment stimulation of macrophages towards an M2 phenotype was clinically predictive of tumor response, we analyzed myeloid phenotypes in the independent validation cohort. We verified that ribociclib resistant tumors had more M2 polarized cells prior to and throughout treatment compared to sensitive tumors (df=6971, t=30.56, p<0.00001) (Figure S7). Across the discovery and validation cohorts, the average M2 differentiation of myeloid cells was significantly higher in ribociclib resistant tumors relative to sensitive tumors prior to and early in treatment (Day 0-14) (df=70, t=2.99, p<0.01) (**Figure 4B, left figure**). Ribociclib sensitive tumors instead had a higher proportion of their myeloid cells in the immune-activating M1 state throughout treatment (df=70, z=2.68, p<0.01) (**Figure 4B, right figure**). The balance of immune-activating versus immune-suppressive myeloid cell frequency was more indicative of response than the total myeloid abundance.

We next measured the heterogeneity in cancer communications that promote macrophage polarization and ribociclib resistance. We first verified the reliability of M2 polarizing communication measurements through known M2 differentiation communication signals including CSF1-CSF1R, ADAM10-AXL and ZP3-MERTK (Figure S8) ^40,41^. We then performed a supervised analysis of the strength of cancer to macrophage signaling across tumors (**Figure 4C**) (Table S21-S24). We identified the range of ligands that cancer cells used to modulate macrophage phenotype and tumor response by selecting communication pathways through which cancer cells: a) signal more strongly with macrophages of resistant than sensitive tumors or b) have stronger cell-cell interactions with M2 versus M1 macrophages. We then compared the strength of communication from cancer to myeloid cells via each M2 differentiation communication pathway in resistant and sensitive tumor samples taken early in each treatment (Day 0-14) (**Figure 4D**). A diverse set of communication pathways were used by cancer cells of ribociclib resistant tumors and which stimulate M2 differentiation, including: CSF1, CLCF1, several FGFs (1/2/7/17/18/23), the TGF*β* family member GDF9, interleukin 5/11/12, MADCAM1 and PLAU (**Figure 4D, right panel**) (Table S21-S22). These ligands have been established to contribute to macrophage M2 polarization and immune suppression via stimulation of macrophage receptors CSF1R, CSF2RB, NRP1 and IL6R ^42–48^. In contrast, macrophages in letrozole resistant tumors did not receive these M2 differentiation communications from cancer cells (**Figure 4D, right panel**) (Table S23-S24). We verified that the cancer cells were the primary contributors of these M2 differentiation communications in ribociclib resistant tumors by comparing the signaling contribution of each non-cancer and cancer cell type (Figure S9). In ribociclib resistant tumors, the cancer cell contribution was 35% greater than the total signal from all cell types in the sensitive tumors.

A comparison of the strength of each M2 differentiation communication across tumors showed that the heterogeneous cancer populations of each resistant tumor used unique combinations of these M2 differentiation communications (Table S21-S22). Additional M2 differentiation communications not identified in the discovery cohort were detected in the validation cohort ribociclib resistant tumors. This diversity indicates that directly blocking all M2 differentiation signals would be challenging. Together these results reveal that M2 macrophage polarization in ribociclib resistant tumors was likely driven by heterogeneous cancer communications from cancer cells that evolved prior to treatment.

### Loss of M1 macrophage cells diminishes interleukin signaling and subsequent CD8+ T cell recruitment and activation

Macrophage polarization to an M1 or M2 phenotype is expected to drive either anti-tumor immune activation or pro-tumor immune suppression, respectively. We therefore examined the communications of macrophages with cytotoxic CD8+ T cells across treatment resistant and sensitive tumors. We first validate the reliability of macrophage to CD8+ T cells communication measurements by testing known signaling effects of these cells and confirmed that individual M1 macrophages sent stronger immune-activating signals (e.g. CXCL9) and M2 macrophages sent stronger immune-suppressing signals (e.g. CXCL13 and CD47) (Figure S10) ^49,50^.

We next determined how the M2 polarization of the entire myeloid population of ribociclib resistant tumors impacted the communication of immune-activating signals to T cells, with an analysis overview presented in **Figure 5A**. This analysis revealed the specific macrophage to T cell inflammatory cytokine communications that diverged during treatment between ribociclib resistant and sensitive tumors. Hierarchical random effects models quantified macrophage to T cell communication differences at end of treatment between resistant and sensitive tumors, while controlling for background patient specific variation in immune state. This revealed that CD8+ T cells of ribociclib resistant tumors received fewer interleukin 15 and 18 (IL-15, IL-18) activation signals from macrophages compared to ribociclib sensitive tumors; with less stimulation of interleukin receptors 2, 15 and 18 on T cells (**Figure 5B**) (Table S25). These receptors are essential for survival, proliferation, and effector differentiation respectively ^51–54^ (IL-15-IL2RA: df=48, t=4.7, IL-15-IL-15RA: p<0.00001, df=27, t=2.33, p<0.05, IL-18-IL-18R1: df=60, t=5.24, p<0.00001). Using the independently profiled validation cohort, we verified that T cells of ribociclib resistant tumors received fewer of each of these IL115/18 activation signals during treatment, while these communications increased in sensitive tumors (Figure S11) (Table S26).

**Figure 5.**
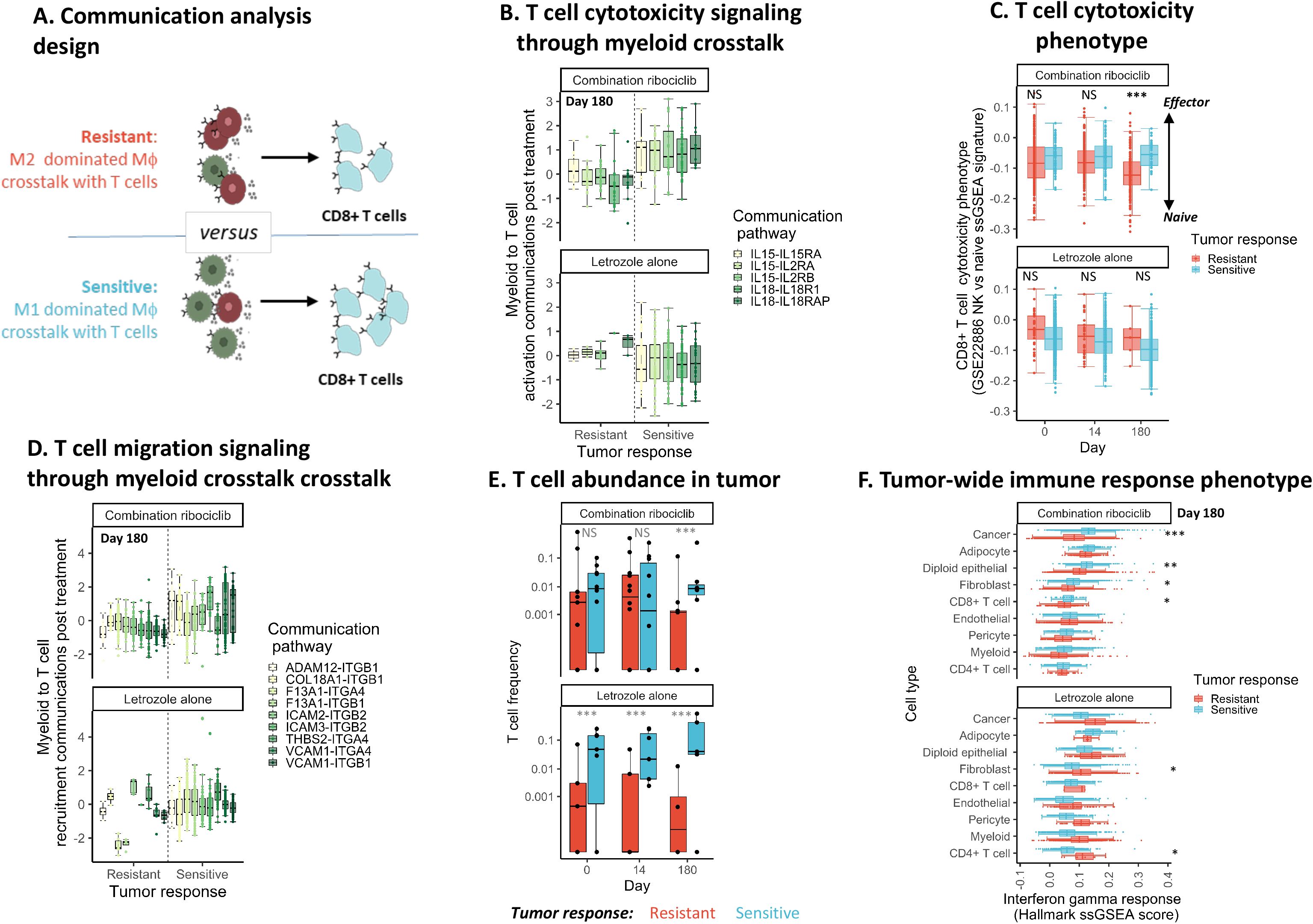
Tumors sensitive to cell cycle therapy maintain CD8+ T cell migration and cytotoxicity through communication with anti-tumor macrophage cells. A) Schematic showing the design of the communications pathway analysis used to reveal the consequence of M2 polarization of the entire myeloid population on the immune activating signals to CD8+ T cells in combination ribociclib resistant tumors. Immune activating inflammatory cytokine communications were selected, using the receptors gene-ontology database signatures. For each immune activating signal, we contrasted the strength of communication sent by the phenotypically diverse myeloid population to CD8+ T cells between treatment resistant and sensitive tumors. We described the strength of communication throughout treatment using a hierarchical regression model and used the Satterthwaite method to calculate t statistics and identify significant divergence in immune activating communication with CD8+ T cells between resistant and sensitive tumors. B) Box and whisker plot showing the weaker CD8+ T cell activating communications received from the myeloid population in combination ribociclib resistant tumors compared to sensitive tumors (top panel right versus left of dashed line) at the end of treatment (Day 180). Colored boxes indicate different CD8+ T cell activating interleukin communication pathways reduced in ribociclib resistant tumors post treatment. C) Box and whisker plot showing the loss of effector differentiation of cytotoxic CD8+ T cells in combination ribociclib resistant tumors at end of treatment. The activation of CD8+ T cell was measured before, during and after treatment with combination ribociclib (top panel) and letrozole alone (bottom panel) using a CD8+ T cell specific ssGSEA pathway contrasting gene expression of naive and cancer killing effector cells (GSE 22886 Naive CD8 T cell vs NK cell up). D) Box and whisker plot showing the weaker CD8+ T cell recruitment communications received from the myeloid population in combination ribociclib resistant tumors compared to sensitive tumors (top panel right versus left of dashed line) at the end of treatment (Day 180). Colored boxes indicate different CD8+ T cell recruiting integrin communication pathways reduced in ribociclib resistant tumors post treatment. E) Box and whisker plot showing the reduction in cytotoxic CD8+ T cell abundance in combination ribociclib resistant tumors at end of treatment. The relative abundance of CD8+ T cells was measured before, during and after treatment with combination ribociclib (top panel) and letrozole alone (bottom panel) and significant differences in abundance between resistant (red) and sensitive (blue) tumors (black points) was determined using logistic regression. F) Box and whisker plot showing the reduced activation of an immune response phenotype across cell type, especially in cancer cells, following treatment in tumors ribociclib resistant tumors. Immune response phenotype measured post-treatment, using the ssGSEA hallmark interferon gamma response pathway score (y axis) of single cells across cell types (x axis) in tumors resistant and sensitive (blue) to combination ribociclib (top panel) and letrozole alone (bottom panel).

We compared this tumor-wide communication from across macrophages with the ability of individual M1 macrophages to crosstalk with CD8+ T cells. We found that significantly fewer T cell activating communications were sent per M1 macrophage in ribociclib resistant tumors, indicating the suppressed activity of cells in this key immune stimulating population (Figure S12). We measured the overall immune activating myeloid to CD8+ T cells communication across immune activating inflammatory cytokine pathways (identified using the gene-ontology database ^55^). This analysis also indicated that the M2 dominated macrophage populations of resistant tumors provided progressively fewer immune activating signals to CD8+ T cells throughout ribociclib treatment (df=185, t=4.48, p<0.0001), whilst immune activation was maintained in sensitive tumors (stronger end of treatment communication vs resistant tumors: df=778, t=6.79, p<0.0001) (Figure S13).

Phenotypic activation of CD8+ T cells was measured using a CD8 T cell specific ssGSEA pathway contrasting gene expression of naive and cancer killing effector cells (GSE22886: “Naive CD8 T cell vs NK cell up” pathway). This analysis showed that activation to an effector CD8+ T cell phenotype was associated with the strength of inflammatory cytokine communications received from macrophages (df=1975, t=7.69, p<0.0001) (Figure S14). In ribociclib resistant tumors, the loss of T cell activating communications was concurrent with the reduction of CD8+ T cell differentiation towards a cytotoxic effector state during treatment (df=786, t=4.98, p<0.0001) (**Figure 5C**).

By the end of treatment, T cells of ribociclib resistant tumors received fewer recruitment signals from macrophages compared to sensitive tumors (Table S25). Stimulation of a variety of T cell integrin receptors was reduced, including ITGB1, ITGB2 and ITGA4 (**Figure 5D**) (ADAM121-ITGB1:df=84, t=5.26, p<0.0001, VCAM1-ITGB2:df=60, t=5.82, p<0.0001, VCAM1-ITGA4:df=117, t=5.94, p<0.0001). These integrins are critical for T cell migration and recruitment through basement membranes ^56,57^. We confirmed that in ribociclib resistant tumors, the decrease of T cell recruitment communications from myeloid cells was linked to a post treatment reduction in T cell abundance (df=21, z=−3.29, p<0.005) (**Figure 5E**). In contrast, a stable abundance of T cells was observed in ribociclib sensitive tumors. Throughout letrozole treatment, T cell abundance was also lower in resistant compared sensitive tumors (df=32, z=33.52, p<0.0001). Again, these results were validated in the independently profiled validation cohort, with T cells of ribociclib resistant tumors receiving fewer recruitment signals than those of sensitive tumors (Figure S15) (Table S26-S28). Together, these results indicate the central role of macrophages in orchestrating T cell activation and recruitment and an effective anti-tumor response during ribociclib treatment.

We then assessed how cancer and non-cancer cells responded to the diverse cytokine communications in the TME. We expected that in immune hot TME’s, the high levels of immune activating signals, and T cell recruitment and activation should induce a phenotypic response across cell types, with activation of interferon regulatory factors (IRFs), induction of IFN-stimulated genes (e.g. interferon gamma-induced proteins) and increased production of antigen presenting major histocompatibility complex molecules (MHC I) allowing recognition and killing of cancer cells ^58,59^. The activation of the interferon gamma response pathway is expected in response to cancer antigens versus interferon alpha response upon viral infection. We measured the interferon gamma response at the end of treatment in cells of each cell type and across tumors, using the Hallmark interferon gamma response ssGSEA signature. The interferon gamma immune response was suppressed across all cell types in ribociclib resistant tumors compared with either ribociclib sensitive tumors (df=19.26, t=−2.23, p<0.05) or tumors treated with letrozole alone (df=21.38, t=−3.29, p<0.005).

Cancer cells in particular exhibited a substantially weaker interferon gamma response post treatment in ribociclib resistant compared to sensitive tumors (**Figure 5F**) (df=788, t=−10.05, p<0.0001). This lack of immune detection was not observed in letrozole resistant tumors (Figure 5E bottom). The result was confirmed in the validation cohort, with resistant cancer cells exhibiting a lower post treatment interferon gamma response than those of sensitive tumors under ribociclib but not letrozole treatment (df=26211, t=−28.06, p<0.0001). We identified communication pathways strongly associated with the cancer interferon gamma response phenotype activation, using Lasso regression (see Methods). Activation of an interferon gamma response in cancer cells was frequently associated with receipt of strong IL-15 signals (28% of tumor subclones) or related cytokine receptors such as: Toll-like receptor 2 (TLR2: 19% subclones), Interleukin 22 Receptor Subunit Alpha 1 (IL22RA1: 17% subclones), Interleukin-12 Receptor Subunit Beta-1 (IL12RB1: 15% subclones) (Figure S16). Together these results indicate that cells of resistant tumors experienced a less hot tumor microenvironment at end of treatment compared to those of sensitive tumors. In particular, cancer cells received fewer immune activating communications that were ultimately linked to a weaker interferon phenotype response to permit immune detection.

We next performed in vitro experiments to examine how CD8+ T cell viability is impacted by a ribociclib treatment and the presence of IL-15 or IL-18 cytokine signals in the environment (**Figure 6A**). The 72-hour T cell viability of two donor populations was reduced by ribociclib treatment (df=31, t=−15.15, p<0.0001) but was much more substantially increased by IL-15 cytokine treatment (df=31, t=24.31, p<0.0001) (but not IL-18 (df=31, t=−0.86, p=0.39)). Furthermore, IL-15 increased T cell viability more under ribociclib treatment than without the drug (df=31, t=3.07, p<0.005). This indicates that IL-15 cytokine treatment can rescue the CD8+ T cell proliferation during ribociclib treatment.

**Figure 6.**
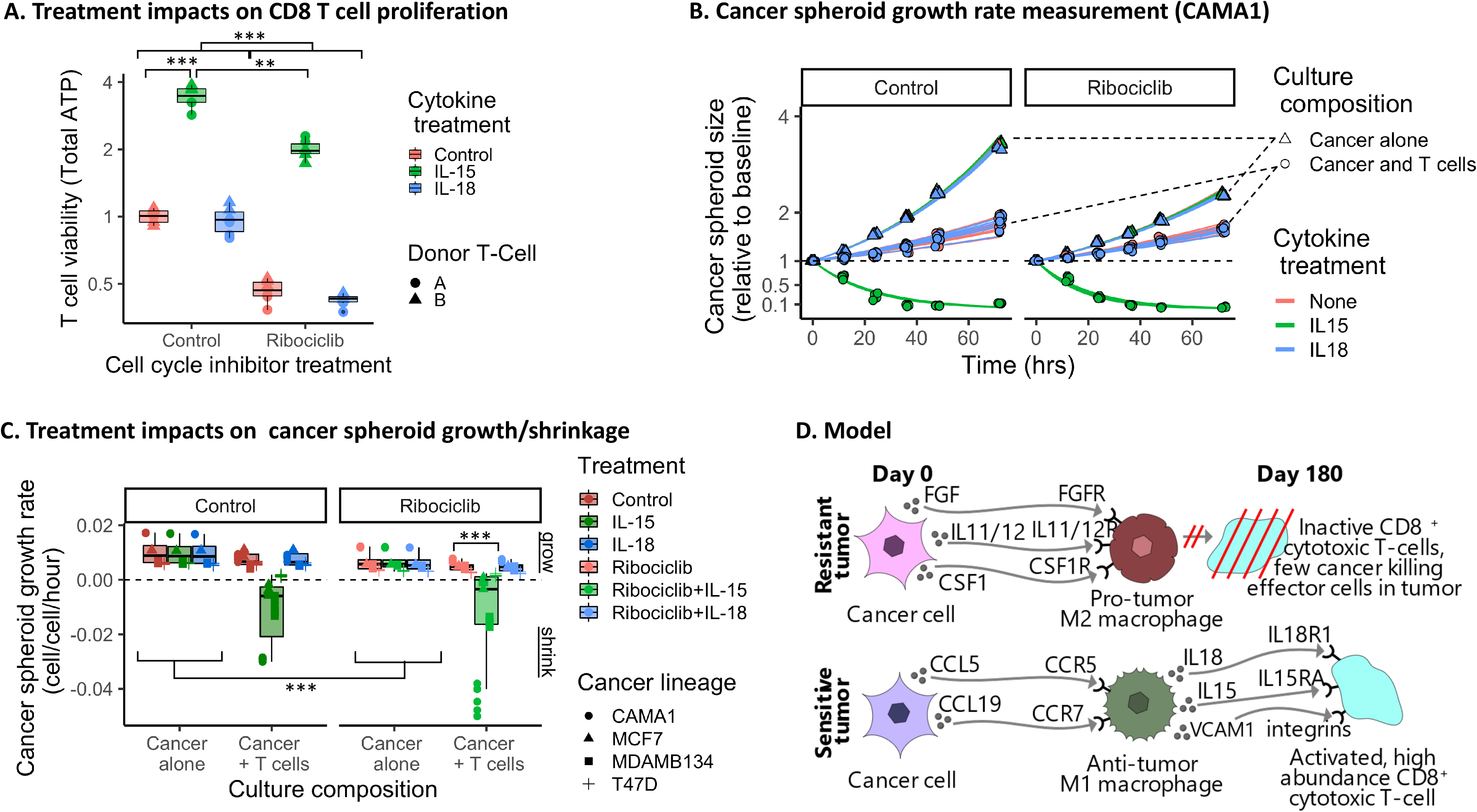
Interleukin 15 boosts CD8+ T cell activation and cancer control even with the anti-proliferative effects of ribociclib. A). Box and whisker plot showing the impact of cell cycle inhibitor (x axis) and cytokine treatment (color) on the viability (total ATP) of CD8+ T cells from two donors patients (point shape). Ribociclib treatment reduced in CD8+ T cell viability and cytokine treatment with IL-15, but not IL-18, substantially boosted CD8+ T cell viability. Under ribociclib treatment, IL-15 provided a significantly greater boost to T cell viability, restoring levels to above the untreated baseline. B). Growth trajectories of cancer spheroids over time in experiments varying cell cycle inhibitor treatment (no drug control vs ribociclib 5um), cytokine treatment (no drug control vs IL-15 vs IL-18) and the presence or absence of CD8+ T cells (present vs absent). For four ER+ breast cancer cell lines (CAMA1, MCF7, MDAMB134 and T47D), cancer spheroid size (point) was measured repeatedly over time using total fluorescence intensity of the lentiviral labeled fluorescent cancer cells. Spheroid size was scaled relative to the initial size following a 1-3 day T cell activation lag. The speed of spheroid growth or decline upon T cell activation was measured by fitting an exponential growth model (colored lines indicate replicate spheroid growth trajectories). The coloration of each replicates’ data points and corresponding fitted line indicates the spheroid growth rate (red= growing, blue= shrinking). C). Box plots showing the shrinkage of cancer spheroids when co-cultured with CD8+ T cells and treated with IL-15, alone or in combination with ribociclib treatment. Cancer spheroid growth rates are compared when cancer cells were cultured alone (left panel) or co-cultured with CD8+ T cells (right panel) under different cell cycle inhibitor treatments (x axis:no drug control vs ribociclib 5um) and cytokine treatment (color: no drug control vs IL-15 vs IL-18). Under IL-15 treatment and when cancer cells were co-cultured with CD8+ T cells, the growth rates of all four ER+ breast cancer cells lines (shapes) were negative (indicating spheroid shrinkage; below horizontal dashed line). This occurred with or without ribociclib treatment (x axis). Replicate experiments using two different donor T cell populations were combined. D). Schematic model summarizing the communication differences of cancer, macrophage and CD8+ T cells between tumors resistant and sensitive to ribociclib treatment. Prior to treatment, resistant tumors have diverse communications from cancer cells that stimulate macrophage polarization to an immune-suppressing M2 phenotype. The lack of pro-immune M1 macrophages diminishes interleukin and integrin signaling and subsequent CD8+ T cell activation and recruitment, preventing effective killing of the quiescent cancer cells.

Next, we examined how ribociclib and the cytokine signals impact CD8+ T cell regulation of cancer population growth. For four cancer cell lines, we compared the growth of cancer mono-culture spheroids with the growth of cancer populations when co-cultured with CD8+ T cells. For each cell line, we assessed how individual or combined ribociclib and IL-15/18 treatments impacted the growth of cancer mono-cultures and co-cultures. **Figure 6B** shows the experimental design and results for CAMA1 cancer cells. The speed of growth or decline of each cancer population was measured by calculating the cancer relative growth rate (rgr) during 75 hour following T cell activation (see methods). The fitted lines show the predicted cancer growth trajectory given the estimated rgr. Finally, we examine the impacts of CD8+ T cells, ribociclib treatment and IL-15/18 signal on cancer growth rates across the four cancer lineages (**Figure 6C**). Ribociclib directly reduced the growth rate of cancer spheroid when grown in monoculture (df=21.6, t=−3.05, p<0.005). The addition of CD8+ T cells without cytokine treatment did not reduce cancer population growth. However, with the addition of IL-15, but not IL-18, CD8+ T cells effectively caused cancer spheroid shrinkage (negative rgr) (df=21.6, t=−7.42, p<0.0001). The efficacy of the IL-15 activated T cells at reducing cancer growth was unaffected by ribociclib treatment (df=21.6, t=−0.01, p=0.99).

Overall, these results indicate that cancer cells of ribociclib resistant tumors avoid immune surveillance through polarization of macrophages to an M2 phenotype, suppressing inflammatory cytokine signaling, T cell activation and recruitment and avoiding cancer cell recognition and killing by immune cells (**Figure 6D**). Interleukin ligands and receptors play a key role in macrophage signaling to activate T cells and the induction of interferon response in cancer cells.

## Discussion

Our analyses of the phenotypes, composition and communication of cell types within the TME revealed the central role of tumor ecosystem-wide signaling in treatment response (Figure 5G bottom). Cancer cells dominate the production of communications within the TME Cancer cells from ribociclib resistant tumors modify the non-cancer behavior of immune regulating myeloid cells, using diverse communications including CSF1, CLCF1, FGF, and TGF family members and interleukin 5/11/12. These resistant tumor specific communications polarize myeloid cells into an M2 phenotype, suppressing subsequent T cell activation and recruitment through loss of interleukin and integrin signaling, respectively. These signaling effects lead to a reduced T cell differentiation and abundances in ribociclib resistant tumors during treatment and to a lack of an interferon response required for cancer recognition and killing by T cells. By connecting communication and phenotypes across cell types, we revealed how communication modulates cell states and tissue function and how treatment targets can be uncovered by studying the corruption and dysregulation of non-cancer cells by cancer. Finally, we revealed that the abundance of immune, stromal and cancer cell types within a tumor provide pre-treatment indicators of patient response to endocrine and cell cycle inhibitor treatments.

### Immunogenicity of ER+ breast cancer

The perspective that breast cancer is immunologically cold and not treatable with immunotherapy is being overturned, yet clinically ER+ tumors are still presumed to be less responsive than other subtypes ^60,61^. However, evidence is accumulating that a strong immune response is indeed essential to ensure tumor response. For example, a high abundance of inactive or regulatory T cells predicts worse overall survival and relapse risk in ER+ tumors ^62^. Conversely, the co-occurrence of many CD8+ cytotoxic T cells with CD4+ helper T cells has been associated with increased progression free and overall survival ^63^. Our results show that within ER+ breast cancers, there exist three distinct archetypical ecosystem compositions: i) a cancer-dominated state, ii) an immune-hot and diverse state, and iii) a fibroblast and endothelial-enriched state. Those initially immune-hot tumors are sensitive to treatment with anti-proliferative endocrine and cell cycle inhibitor treatments. This result suggests that preemptively or concurrently heating cold tumors may improve response to cell cycle and endocrine inhibition.

### Immunotherapy for ER+ breast cancer

Immunotherapy strategies have been developed to treat immune hot and cold tumors ^64^, however they are not currently used based on a mechanistic single cell perspective of treatment resistance or tailored to specific types of immune-related signaling. Our results highlight the treatment dependent role of the TME in predisposing tumor resistance. Immune activation at baseline and during treatment did correlate to improved tumor response to both letrozole alone and combination ribociclib treatments. However, response to ribociclib was far more dependent on the composition, phenotypes and communication of immune cells, particularly macrophages, than the letrozole response. This may be due to ribociclib’s effect on immune cells, as it is known that cell cycle inhibitors can cause low white blood cell counts and block T cell proliferation ^65–67^. Thus, the dual impact of ribociclib on both cancer cell and immune cell proliferation and activity could impact a tumor’s response. One hypothesis is that recruitment, abundance, and activation of cytotoxic T cells, through IL-15/18 and other cytokine signaling, impacts how tumors respond to therapy. It may be insufficient to block cancer cell growth alone; immune cell recognition, cytotoxic effects, and clearance of cancer cells may be a critical component of tumor response to cell cycle therapies. In contrast, the targeted effect of letrozole on the cancer cell specific estrogen growth factor pathway have less impact on immune cell signaling and T cell cytotoxicity.

We propose that a durable tumor response to cell cycle treatment requires the killing of the quiescent cancer cells by the immune system. Otherwise, the cancer cells eventually evolve to bypass the effects of anti-proliferative drugs or manipulate the TME to engineer a more proliferative microenvironment through stromal interactions and growth factor secretion. We have previously shown that tumor phenotypic evolution during combination ribociclib therapy leads to the emergence of cell cycle reactivation through a shift from estrogen to alternative growth signal-mediated proliferation in this early-stage ER+ breast cancer population ^10^. Here we show how the pre-treatment corruption of cancer to macrophage communications drive suppression of immune function, enabling cancer cells to survive ribociclib treatment. These data identify several immunotherapeutic targets to overcome cell cycle inhibitor resistance: i) block the macrophage polarizing communications sent by cancer cells, ii) enhance M1 differentiation and iii) directly activate cytokine signaling to recruit and activate effector T cells and promote antigen presentation.

### Blocking diverse cancer-myeloid communication

The diversity of cancer communications used concurrently to induce the same phenotypic shift of macrophages towards a pro-tumor state presents a therapeutic challenge. Treatments aiming to block the cancer-macrophage crosstalk would need to be broad and would thus likely incur toxicity or tolerability problems. Drugs targeting specific corrupting cancer communications (e.g. an CSFRi such as ARRY-382) would be unlikely to block all communication pathways inducing the M2 transition in a specific patient’s tumor. This suggests that a better treatment approach would be to target the downstream consequences on macrophage phenotype, their communications and impacts on T cell function.

### Enhancing M1 differentiation

The switching of macrophage polarization from an M1-M2 phenotype is known to regulate pro-versus anti-tumor immunity ^68^. Our results uncover the central role of macrophage M2 polarization in predisposing resistance to cell cycle inhibition. Drug sensitivity was higher in tumors with more abundant dendritic cells and more inflammatory M1 macrophages. Several promising immunotherapeutic strategies are emerging to promote M1 differentiation ^69^. These include: local low-dose irradiation ^70^, intra-tumoral IL-21 injections ^71^, immune checkpoint inhibitors ^72^, autologous GM-CSF vaccines ^73^ or chimeric antigen receptor macrophage transfer ^74^.

### Activating cytokine signaling in ER+ tumors

Providing the signals to promote CD8+ cytotoxic T cell activation, recruitment and proliferation is a more direct method to overcome ribociclib resistance, without treating the cause of immune suppression. Our analyses identified that sensitive tumors had robust IL-15 signaling between macrophages, T cells and cancer cells, leading to increased T cell recruitment, activation and stronger antigenic interferon response of cancer cells. In both breast and colon cancer murine models, IL-15 promotes tumor destruction and reduces metastasis through T cell activation ^75,76^. Furthermore, IL-15 agonists can overcome the immunosuppressive effects of antiproliferative drugs (e.g. MEK inhibitors) ^77^. Consequently, IL-15 is recognized as a highly promising immunotherapeutic agent, with clinical studies underway testing efficacy as an adjuvant or combination treatment for metastatic solid tumors ^78^. Benefits over other FDA approved cytokines such as IL2 include reduced toxicity and avoidance of Treg differentiation and activation-induced CD8+ T cell death ^79^. Superagonists have been developed to enhance bioactivity and stability, prolonging antitumor immunity. A major current obstacle is identifying the tumor types to target with IL-15 along with the optimal dose and timing. Our results show that the use of single cell transcriptomic profiling of the TME can identify tumors to target by uncovering the strength of inflammatory communication and the activation of cytotoxic T cells.

Our approach of serially profiling the cancer and non-cancer cells across a cohort of patient tumors both resistant and sensitive to a given therapy allows identification of the key TME driven mechanisms of resistance. Insights into the tumor composition, communication and phenotypes can guide individualized treatment strategies. The TME’s composition can predict tumor sensitivity to cell cycle and endocrine therapies, communications show the mechanism of dysregulation and reveal personalized treatment targets and the phenotype of cells provide early indicators of treatment success.

## Methods

### Patient cohort and sample collection

Patient tumor core biopsies were collected prospectively under Clinical Trial #NCT02712723 ^21^, during a randomized, placebo controlled, multicenter investigator-initiated trial led by Dr. Qamar Khan at the University of Kansas Medical Center (IND #127673). The trial entitled FELINE studied Femara (letrozole) plus ribociclib (LEE011) or placebo as neo-adjuvant endocrine therapy for women with ER-positive, HER2-negative early breast cancer. Postmenopausal women with pathologically confirmed non-metastatic, operable, invasive breast cancer and clinical tumor size of at least 2 cm were enrolled from 10 centers across the United States. Invasive breast cancer had to be ER positive (≥66% of the cells positive or ER Allred score 6–8) and HER2 negative by ASCO-CAP guidelines.

One hundred and twenty patients were randomized equally across three treatment arms (40:40:40). Arm A received letrozole plus placebo, Arm B letrozole plus ribociclib 600 mg daily for 21 out of 28 days of each cycle and Arm C received letrozole plus ribociclib 400 mg continuously. Protocol therapy was continued until the day before surgery. Tumor response to treatment was assessed using multiple imaging modalities. Mammogram, MRI and ultrasound of the affected breast were performed at baseline and a mammogram and ultrasound was performed at completion of neoadjuvant therapy. MRI of the breast was performed after completion of 2 cycles of treatment (Day 1 of cycle 3). Serial tissue biopsies using a 14-gauge needle were mandatory, providing three core tumor sample over the course of treatment: baseline (Day 0), Cycle 1 Day 14 (Day 14), and end of treatment (Day 180). Immediately after collection, biopsy samples were snap frozen embedded in optimal cutting temperature (OCT). Informed consent was obtained from all patients following protocols approved by the institutional IRBs and in accordance with the Declaration of Helsinki. The study was approved by University of Kansas Institutional Review Board (protocol #CLEE011XUS10T).

### Single nuclei RNA sequencing and processing

Tumor single cell nuclei were isolated from OCT embedded core tumor biopsies using a modified lysis buffer containing 0.2% Igepal CA-630 as previously described ^*80*^. Single cell RNA-Sequencing (scRNAseq) was performed on single nuclei suspensions using 10X Genomics Chromium platform as previously described ^10^. Sequence reads were processed with BETSY and CellRanger v3.0.2, which aligned reads to reference genome (GRChg38) using STAR v2.6.0 ^81^. For each sample, a gene-barcode count matrix was generated containing counts of unique molecular identifiers (UMIs) for each gene in each cell.

We reanalyzed the validation cohort cells to recover intermediate-quality non-cancer cells that were excluded based on the filters used originally in the discovery cohort analysis. Cells were clustered based on the percentage of mitochondrial genes using k-means clustering (k =4). We filtered out high mitochondrial content clusters (centroids = 55 and 87% mitochondrial genes) and retained low percent mitochondrial genes (centers = 0.2 and 20%). We further filtered out cells classified as epithelial by SingleR analysis^23^ and with less than 100 genes expressed.

### Cell type classification and verification

We obtained transcriptional profiles of 424,581 single cells, using stringent quality controls to ensure high-coverage, low mitochondrial content, and doublet removal (as described in ^10^). Broad cell types were annotated using singleR^23^, cancer cells were identified by their frequent and pronounced copy numbers amplification using inferCNV^24^. Cell type annotations were verified by cell type specific marker gene expression and Umap/TSNE analyses ^10,25^. Granular immune subtype annotations were obtained using our recently published ImmClassifier machine learning method, which has been validated by flow cytometry comparisons ^26^.

### Machine learning classifier for cell type annotations

Cell subtype annotations for the discovery and validation cohorts were consistently annotated by training a random forest machine learning classifier to identify cell types using the well-curated discovery cohort data. We then applied the classifier to predict cell type annotations in the validation cohort. First, we identified the marker genes associated with each cell type in the discovery cohort cells, using a negative binomial test to find genes differentially expressed in each cell type relative to all others. Genes expressed in > 25% of cells in at least one group and showing a log fold change in expression > 0.25 between the groups were selected as candidate markers. We additionally included cell cycle score (G2M and S scores calculated by Seurat’s CellCycleScoring function) as latent variables. The top 100 marker genes of each cell type were selected as candidate features in the machine learning analysis. The classifier was constructed using SingleCellNet, a top pair random forest approach to predict each cell type^82^. First, we split the high-quality C1 cohort into a training and validation subset. We then trained the classifier on the training subset with the parameters nTopGenes = 25, nRand = 100, nTrees = 1000 and nTopGenePairs = 50. Performance assessment in the held-out validation subset showed good performance with area under the receiver operator curve > 0.9 (Figure S1). To confirm the consistency of the discovery and validation cohorts, all cells were projected into a common UMAP space, using the first 10 principal components of the scaled expression levels of 100 marker genes associated with each cell type (Figure S1). We verified that the UMAP clusters, indicating a major biological cell type, were assigned consistent cell type annotations using the across cohorts using the manual curation and machine learning classification approaches. Most cell type were uniquely assigned to a single cluster and this accuracy was further improved by retaining cells with a cell type prediction probability > 0.75.

### Archetypal tumor compositions

The composition of each tumor sample was summarized by calculating the proportion of each cell subtype to correct for sampling variation. The compositional similarity of each tumor samples was measured using the pairwise Manhattan distance. This quantified the fraction of each tumor composition that would need to be altered to generate the compositional profile of each other sample. Compositionally similar tumors and collections of cell types with correlated abundances were grouped using hierarchical clustering (method= ‘ward.D2’).

To identify archetypal ER+ breast cancer tumor compositions, we projected all tumor samples into a composition landscape, using UMAP to account for the non-linearity and non-normality of compositional data ^25^. Highly compositionally similar tumors located close together in this ordination space and distant from tumors with divergent compositions. We then applied a Gaussian Mixture model (GMM) and Bayesian information criterion to probabilistically identify distinctly similar clusters of tumor samples ^83^. This identified the appropriate number of archetypal tumor compositions supported by the data and classified each tumor sample into an archetype. Major compositional differences between archetypal compositions were identified using Dirichlet regression (R package ‘DirichletReg’ v0.7-1) and the rank correlation of UMAP TME composition axes with cell type frequencies.

### Subclonal cancer composition and evolution from scRNA copy number alteration

Cancer subclonal populations of each tumor sample were identified through ‘infercnv’ analysis (R package ‘infercnv’ v1.0.2; cutoff=0, min_cells_per_gene=100 or 500, cluster_by_groups=T, HMM=T, analysis_mode=“subclusters”). Genomic regions of copy number alteration in each cell were detected relative a subset of 500 reference immune or stromal cells, using the count matrix. Then, cancer populations with distinct copy number profiles were defined as cancer subclones of a patient tumor using hierarchical clustering (R package ‘fastcluster’ v1.1.25; method=‘ward.D2’)^84^. Clusters with distinct copy number profiles were defined as subclones for each patient. Single-cell grouping was performed based on hierarchical cluster analysis.

### Cell phenotypes from Gene Set enrichment analysis

The gene expression count matrix of each cell type was filtered to keep genes expressed in at least 10 cells, zinbwave normalized with total number of counts, gene length and GC-content as covariates (R package ‘zinbwave’ v 1.8.0; K=2, X=“~log (total number of counts)”, V=“~ GC-content + log (gene length)”, epsilon=1000, normalizedValues=TRUE) ^85^. Single sample Gene Set Enrichment Analysis (ssGSEA) scores of 50 hallmark signatures (MSigDB, hallmark) and 4725 curated pathway signatures (MSigDB, c2) were calculated for each cell using the normalized count matrix in GSVA (R package ‘GSVA’ v1.30.0; kcdf=“Gaussian”, method=‘ssgsea’) ^86,87^.

### Communication from across diverse cell type populations through Tumor-wide Integration of Signaling to Each Receiver (TWISTER)

To uncover networks of communication from across diverse cell type populations and received by individual cancer and non-cancer cells of a tumor, we developed the TWISTER algorithm. This measures population level communication using single-cell gene expression (count per million). We first extended individual level cell-cell interaction (CCI) approaches (reviewed in ^12^) to measure communications received from entire cell type populations or from across the entire tumor population (tumor-wide communication), accounting for tumor composition and within cell type phenotypic heterogeneity. The tumor-wide communication metric was derived by formulating a differential equation model of tumor ecosystem signaling. This describes the change in concentration of a signaling ligand molecule (*S*) in the TME as following:

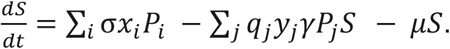

Signals are produced by cells in the TME at a rate proportional to their expression of the signaling ligands. Within cancer and non-cancer cell types, subpopulations vary phenotypically and differ in ligand gene expression, with subpopulation *i* having ligand expression *x_i_*. Ligands produced by a cell are released into TME at rate *σ*. The total signal production by each cell subpopulation is proportional their abundance in the TME (*P_i_*) and the total signal production across the TME is given by the sum of production across all cell subpopulations. Signaling ligands are removed from the TME through decays or diffusion at rate (*μ*) or when bound to a receptor on a receiving cell (receptor binding rate=*γ*) and taken up. Phenotypically different subpopulations within cancer and non-cancer cell types have differing receptor concentrations, with receptor density of cell type j depending on its receptor gene expression (*y_j_*). The total ligand uptake by each cell subpopulation is proportional to their abundance in the TME (*p_j_*) and the fraction of molecules are taken up and removed from the TME once receptor bound. The total signal uptake across is given by the sum of uptake across all cell subpopulations.

The steady state analysis of the TME signal concentration is given by:

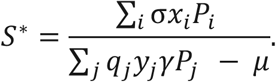

Assuming ligand release after receptor binding (*q_j_* is small), the strength of signal transmitted from all cell subpopulation in the TME (tumor-wide communication) to a focal receiver cell in subpopulation j is given by:

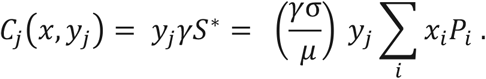

Given a sampled tumor composition 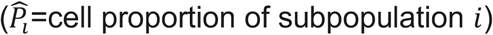, the tumor-wide communication transmitted via ligand-receptor pathway *k* to a receiver cell (of type *j*) can be measured given a vector of ligand gene expression for each cell types present (*x_k_*), and the receptor expression of the receiving cell (*y_jk_*) as: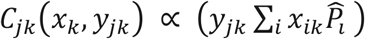 **(Figure 1C)**.

This generalizes the CCI approaches using the ligand-receptor product and expression correlation method (e.g. CCCExplorer, ICELLNET, NATMI, NicheNet and scTensor; reviewed in ^12^) to the broader tumor ecosystem perspective. Rather than measuring one-one interactions between individual cells, *C_jk_*(*x_k_, y_jk_*) measures the many-one communication strength a focal cell receives from the diversity of cells that are releasing communications into the TME. This is again distinct from the many-many mapping of communication implemented to quantify the probability of cell-cell communication between two cell types (e.g. in CellChat)^14^. TWISTER therefore allows an assessment of how phenotypically diverse populations of cells contribute communications to the signaling reservoir in the TME to stimulate the receiver cell, accounting for the abundance and ligand production of each signaling phenotype **(Figure 1C)**.

We validated that by restricting tumor-wide communications to individual level communications between one sending cell of one cell type and another receiving cell, measurements are consistent with individual level cell-cell interactions obtained using the ligand-receptor correlation/expression product method (as used in CCCExplorer, ICELLNET, NATMI, NicheNet and scTensor; reviewed in: ^12^). Our model also shows how tumor-wide communications generalize the established CCI approach to the broader tumor ecosystem perspective. Crucially, instead of just revealing how an individual cell of one cell type communicates with a cell of another type (individual one-one cell crosstalk), TWISTER allows an assessment of how a phenotypically diverse population of cells within each major cell type (e.g. macrophages in distinct states) contribute communications to the receiver, accounting for the abundance and ligand production of each signaling phenotype (population many-one cell crosstalk). This is distinct from methods such as CellChat which us cell counts to weight the probability that an individual of the two cell types interact (i.e. frequency of individual one-one cell crosstalk).

### Measuring tumor-wide communications received from diverse cell phenotypes

We apply TWISTER to measure tumor-wide signaling from diverse non-cancer cell sub-populations and heterogeneous cancer lineages to receiving cells. We first resolved diverse subpopulations of each cancer and non-cancer cell type (e.g. macrophages in different differentiation states). For each broad cell type we generated a cell-type specific UMAP based on ssGSEA profiles, with the intrinsic UMAP dimensionality determined using the packing number estimator^88^. We then break down each cell type into subtypes of at least 30 cells with coherent phenotypes and of equal interval width along each phenotype axis. This allowed cell types with relatively continuous phenotypic variation, such as macrophages, to be subdivided into an ordered set of cell states along multiple axes of phenotypic heterogeneity and maintains phenotype covariance structure. We then calculated the relative abundance of each subpopulation of each major cell type within a tumor sample 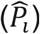.

We next used a curated LR communication database ^89^ to define a set of 1444 LR communication pathways (*C_jk_*(*x_k_, y_jk_*)) based on known protein-protein interactions. We extracted single-cell expression of a ligand and used mean CPM of a cell subpopulation as a metric of signal production (*x_k_*) and the mean CPM receptor expression to quantify signal receipt by a focal cell (*y_jk_*). We calculated activity of each LR communication pathway (*k*=1:1444) between each pair of sending (*i*) and receiving (*j*) subpopulations (*i* → *j*) within a tumor: 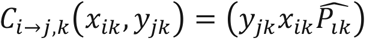

### Strength of communication between each cell type: contribution and receipt of signals

To obtain communications via communication pathway *k* between broad cell types, we totaled signals from sending cell type populations (1: *n* ligand producing subpopulations of a cell type) and averaged signals to receiving cells (across 1: *m* signal receiving subpopulations of a cell type). We use a weighted average so that the signal to each receiving cell type population is weighted by abundance (**Figure 1x**):

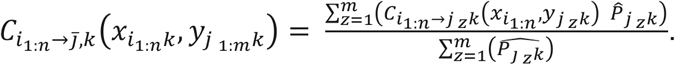

We refer to this as the strength of communication from a cell type population to a typical cell of another type. For example, the contribution of *n* heterogeneous cancer cell populations (*Cancer*_1:*n*_) to the communication with a myeloid cell of phenotype *z* via ligand *x* and receptor *y* is given by:

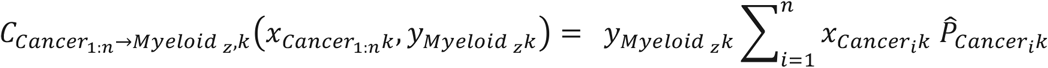

The strength of communication received by a typical myeloid cell in a sample 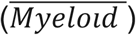 from the diversity of cancer cells is given by:

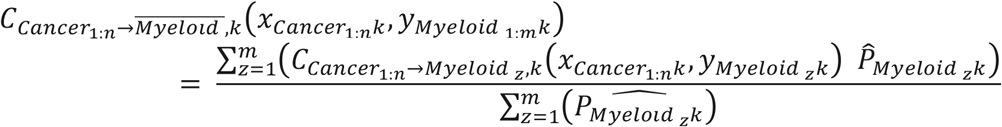

This was repeated for each LR communication pathway between cell types and across tumor samples. Each communication pathway has a distinct potency to modulate cellular phenotype and behavior and so communication pathway scores were standardized (mean=0, sd=1) across patients, preventing highly expressed ligand-receptor pairs dominating communications. The average strength of communication from one cell type to another across LR pathways 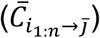 was measured by the median standardized communication from one cell to another.

### Validation of communication measurements

We validated the method in peripheral blood immune cells, in which communications between cell types are well known and distinctly different from those expected in tumor biopsy samples ^90^. Our approach successfully recovered the expected communication network, with myeloid cells having a central role in communicating via cytokine pathways with many cell types (Figure S17). We also validated that we can recover canonical cell type specific communications including receipt of: macrophage colony stimulating factor primarily in myeloid cells, vascular endothelial growth factor (VEGF) in endothelial cell, fibroblast growth factor (FGF) in fibroblasts, epidermal growth factor (EGF) in epithelial cells and C-C chemokine receptor type 5 (CCR5) in T cells (Figure S3). Finally, we compared the measured differences in communication between the resistant and sensitive tumors of the discovery and validation cohort. This verified the high degree of consistency in the signals each cell type received via each LR communication pathway tumor response groups across the two cohorts (*R*^2^ = 0.81) (Figure S18).

### Cell type communication differences between resistant and sensitive tumors: Bootstrapping randomization

Contrasting communication across many biopsies, rather than within individual samples, provided comparative insights into the evolution of communication during treatment and the cell type communications that distinguish resistant and sensitive tumors. We contrasted the networks of communication between cell types in resistant and sensitive tumors and examined how communications changed throughout treatment with ribociclib or letrozole. We determined the difference in the average strength of communication from one cell type to another 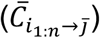 between resistant and sensitive tumors. To identify which cell type communications significantly differed between tumors resistant and sensitive to each treatment, we perform a bootstrapping randomization analysis. We repeatedly shuffled the observed cell type L-R communications across resistant and sensitive tumors to remove any response related structure of the communication network. For 1000 randomized communication networks, the difference in average communication was recalculated. The distribution of communication differences produced by chance in the randomized networks (null model: average communication does not differ between resistant and sensitive tumors) was then compared to the observed difference in the average communication between cell types.

Using the mean and standard deviation of the communication differences in the randomized networks, z statistics were calculated to indicate how much each cell type’s communication differed between resistant and sensitive tumors. Randomization p-values were calculated by the rank of the observed average communication difference within the distribution of randomized differences between resistant and sensitive communication networks. A Holm’s conservative correction for statistical significance was applied to correct for multiple comparisons.

### Diverging communication networks between cell types in resistant and sensitive tumors

We obtained the expected cell type communication of one cell type to another in resistant and sensitive tumors at each time point of each treatment. This summarized the average strength of communication 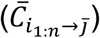 of each individual tumor within each response category and treatment time point. We then calculated the proportional change in expected cell type communication in resistant and sensitive tumors, relative to the baseline overall average. Post treatment cell type communication networks were then described by directed weighted network graphs, constructed for resistant and sensitive tumors. Cell types are represented by a network nodes and the proportional changes in communication are described by the weight of the vertex from one cell type to another (indicated by arrow width).

### Communication pathway analysis: identifying response related communications

For each LR communication pathway, we contrasted the strength of communication between cell types in tumors resistant versus sensitive to each treatment. We used log-linear regression to describe trends in cell type communication within resistant and sensitive tumors 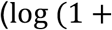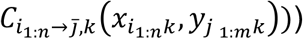. General communication trends during treatment and changes specific to resistant tumors were detected using likelihood ratio tests. Significant differences in the strength of cell type communication between resistant and sensitive tumors either before or after treatment were identified by using ANOVA on the endpoint data (Day 0 and 180 separately). We accounted for multiple comparisons using false discovery rate (FDR) p-value correction. To identify broadly divergent communications between resistant and sensitive tumors before treatment, we enumerated the detected communications sent and received by each cell type.

### Myeloid phenotype reconstruction

The diversity of myeloid phenotypes was examined through UMAP analysis of single cell transcriptional profiles (log(1+CPM)). Genes with greater than 5% coverage in cells were used. Dendritic cell and macrophage cell subtype annotations obtained from ImmClassifier ^26^ were overlaid onto the UMAP, confirming consistent identification of distinct cell population between approaches. The M1-M2 phenotype gradient across the UMAP (dimension 2) was identified using the rank correlation of UMAP axes each genes expression. To test for pre-treatment differences in macrophage polarization between resistant and sensitive tumors, we fitted a hierarchical linear model describing the M1-M2 phenotype score differed in myeloid cells from resistant and sensitive tumors, accounting for patient specific heterogeneity in myeloid phenotype and the shared TME of cells within a sample. Finally, M1 macrophages were defined as having less average M1-M2 phenotype scores and other macrophages classified into the M2 phenotype.

### Verifying that myeloid polarization predicts resistance

We next confirmed that ribociclib resistant tumors can be identified early in treatment (Day 0-14) by the increased M2 differentiation of their myeloid cells. We contrasted the phenotypes of myeloid cells in resistant and sensitive tumors in the independently profiled verification cohort. We characterized the phenotypic heterogeneity of myeloid cells in the verification cohort. We used the fitted UMAP model, trained with the discover cohort CPM data, to project myeloid cells of the validation cohort into a consistent myeloid phenotype space. This provided equivalent M1-M2 phenotype scores for each myeloid cell of the validation cohort.

We then assessed whether myeloid cells of ribociclib resistant tumors showed greater M2 differentiation in the validation cohort. We applied three complementary analyses. First, we applied a hierarchical linear model to test for differences in single cell M1-M2 phenotype scores between resistant and sensitive tumors throughout treatment, accounting for patient specific heterogeneity in myeloid phenotype. Second, we summarized the mean M1-M2 phenotype score of myeloid cells in each tumor. We then contrasted the mean myeloid differentiation early in treatment (Day 0-14) between resistant and sensitive tumors using ANOVA. Finally, we categorized cells into M1 and M2 phenotypes and compared the relative abundance of M1 and M2 macrophages resistant and sensitive tumors early in treatment, using logistic regression to describe the proportion of M1 cells in a tumor biopsy using treatment and resistance outcome as predictors.

### Measuring targeted cancer cell signaling to M1 and M2 macrophages

We identified cancer cell communications that predominantly target either M1 or M2 macrophages. First, we measured cell-cell interactions between phenotypically diverse cancer and macrophage subpopulations in each tumor sample, using the ligand-receptor product approach ^12^. For each tumor sample, we calculated the average cell-cell interaction of cancer cells with each macrophage via each communication pathway. We contrasted the log cancer-macrophage cell-cell interaction received by M1 and M2 macrophages, using a hierarchical linear model to detect differential communication with myeloid cell types and to account for baseline tumor specific differences cancer-macrophage communication. Significant differences in cancer communication with M1 and M2 cells were identified using likelihood ratio tests contrasting: i) the full model with communication to M1 and M2 cells differing and ii) the nested null model with no difference in communication. The twenty most significantly activated communications with M1 and M2 macrophage were assessed.

### Contrasting the heterogeneity of cancer to macrophage communications across resistant and sensitive tumors

We combined the list of communication pathways through which cancer cells: a) communicate more strongly with macrophages in resistant than sensitive tumors (see Communication pathway analysis) and b) have stronger cell-cell interactions with M2 versus M1 macrophages (as described above). From this list, we identified the ligands the cancer cells used to modulate macrophage phenotype and tumor response. Communication pathways binding these ligands were defined as M2 differentiation communications and selected for supervised analysis. We contrasted the strength of communication from cancer to myeloid cells via each M2 differentiation communication pathway in resistant and sensitive tumors samples taken early in each treatment (Day 0 and 14). Heatmaps were used to visualize the heterogeneity of communication pathway activity across tumors.

### Cancer and non-cancer cell type contributions to myeloid polarizing communications

We next determined which cell types most strongly contributed to myeloid differentiation communications. We extracted the strengths of each M2 differentiation communication sent from each cell type to myeloid cells. For each tumor sample, the median standardized M2 differentiation communication from each cell type was calculated. We then contrasted the average M2 differentiation communication sent by each cell type in resistant and sensitive tumors and under each treatment.

### Identifying differential communications of M1 and M2 myeloid cells with CD8+ T cells

We next identified myeloid communications with T cells primarily produced by either M1 and M2 myeloid cells. We first measured cell-cell interactions between phenotypically diverse macrophage and T cell subpopulations in each tumor sample, using the ligand-receptor product approach ^12^. For each tumor sample, we calculated the average cell-cell interaction of M1 and M2 myeloid cells with T cells via each communication pathway.

We then identified communication pathways by which T cells received significantly different cell-cell interactions from M1 and M2 myeloid cells. The log macrophage-T cell interactions from M1 and M2 macrophages were contrasted, using hierarchical linear models. A patient specific random component accounted for the heterogeneity in immune communication between TME’s and a random component associated with T cell phenotype accounted for the diversity of T cell activation phenotypes within and between tumors. Significant differences in T cell communication from M1 and M2 cells were identified using likelihood ratio tests contrasting: i) the full model with communication from M1 and M2 cells differing and ii) the nested null model with no difference in communication.

### Contrasting M1 communication with T cells in resistant and sensitive tumors

Next we isolated the M1 polarized macrophages and examined their communication with T cells in resistant and sensitive tumors. For each communication pathway, we contrasted M1 macrophage to T cell interactions (log transformed) in resistant and sensitive tumor using hierarchical linear models. Again a patient specific random component accounted for the heterogeneity in immune communication between TME’s and a random component associated with T cell phenotype accounted for the diversity of T cell activation phenotypes within and between tumors. A likelihood ratio test was used to detect communication pathways significantly differing between resistant and sensitive tumors.

### Diverging inflammatory communication from myeloid cells to CD8+ T cells in resistant and sensitive tumors

We next determined how the M2 polarization of the myeloid population in ribociclib resistant tumors impacted the communication of immune cytokine signals to T cells. First the subset of immune activating inflammatory cytokine communications were identified, using the receptors gene-ontology database signatures ^55^. For each CD8+ T cells we totaled the signal received from all myeloid subpopulations within that tumor sample via each inflammatory cytokine communication pathway. To obtain the overall immune activating communication received by each CD8+ T cell from the myeloid population, we averaged across communications pathway scores after scaling and log transformation.

We then analyzed at the single cell level how each of the immune activating communications from myeloid to CD8+ T cells diverged during treatment in resistant and sensitive tumors. Using a hierarchical regression model, we described pre-treatment differences in CD8+ T cell activating communication between resistant and sensitive tumors and temporal change during treatment (as previously described in ^10^). Significant divergence in immune activating communication with T cells of resistant and sensitive tumors was determined using a two-tailed t-test. The Satterthwaite method was applied to perform degree of freedom, t-statistic and p-value calculations, using the ‘lmerTest’ R package ^91^.

### Linking myeloid inflammatory communications to CD8+ T cell activation

We characterized how the differentiation and activation to an effector CD8+T cell phenotype was related to the strength of immune cytokine communication they received from myeloid cells. For each CD8+ T cell, differentiation was measured using a CD8+ T cell specific ssGSEA pathway contrasting gene expression of naive and cancer killing effector cells (GSE 22886 Naive CD8 T cell vs NK cell up). The single cell activation state was then linked to the immune activating communication received from across the myeloid population (measured above). Each CD8+ T cells differentiation state was then linked to the inflammatory cytokine communication it received from across the myeloid population. Linear regression was used to measure the increase in T cell activation with increasing inflammatory communication. The strength of inflammatory cytokine communication received was discretized into deciles of signal strength and the distribution of phenotypic state assessed in cells receiving each level of stimulus.

We then analyzed at the single cell level how the CD8+ T cell activation diverged during treatment in resistant and sensitive tumors. Using a hierarchical regression model, we described pre-treatment differences in CD8+ T cell activation between resistant and sensitive tumors and temporal change during treatment (detailed in ^10^). Significant divergence of CD8+ T cell activation in resistant and sensitive tumors was determined using a two-tailed t-test. The Satterthwaite method was applied to perform degree of freedom, t-statistic and p-value calculations, using the ^91^.

### Contrasting T cells relative abundance during treatment in resistant and sensitive tumors

Differences in T cell abundance between resistant and sensitive tumors were analyzed at each treatment time point and separately for tumors receiving each treatment, using logistic regression. We identified significant differences in the proportion of T cells between tumor response groups using a two-tailed Wald-test to generate z statistics and p values.

### Comparing post treatment immune response across cell types in resistant and sensitive tumors

We compared the difference in immune response observed across all cell types between tumors resistant and sensitive to each treatment. For each single cell observed at the end of treatment, we measured immune stimulation using Hallmark Interferon Gamma response ssGSEA pathways.

We analyzed the difference in Interferon Gamma response between treatment resistant and sensitive tumors, using a nested hierarchical regression model to account for the patient specific differences in immune response and the between cell type differences in this phenotype. Cell type specific random effects were nested within the patient random component, reflecting the occurrence of each cell type within different patient tumors. Significant divergence in Interferon Gamma response between resistant and sensitive tumors treated with combination ribociclib or letrozole alone were determined using two-tailed t-tests.

We then analyzed at the single cell level how the CD8+ T cell activation diverged during treatment in resistant and sensitive tumors. Using a hierarchical regression model, we described pre-treatment differences in CD8+ T cell activation between resistant and sensitive tumors and temporal change during treatment (described in ^10^). Significant divergence of CD8+ T cell activation in resistant and sensitive tumors was determined using a two-tailed t-test, again using Satterthwaite method.

### Linking inflammatory communications to cancer interferon gamma response

We next determined the how cancer cell phenotypes responded to increasing inflammatory cytokine communications in the TME. We assessed the cancer cells interferon response phenotype, using their hallmark interferon gamma response ssGSEA scores. This cancer phenotype measured intracellular transduction of cytokine signals to the nucleus, induction of interferon regulatory factors (IRFs), IFN-stimulated gene activation (e.g. Interferon gamma-induced proteins) and the production of antigen presenting major histocompatibility complex molecules (MHC I) allowing recognition and killing of cancer cells ^58,59^.

To determine the major communication pathways stimulating a cancer cell interferon gamma response, we next calculated the strength of the communication each cancer cell received from across the TME via each ligand receptor pathway. For each cancer cell, we coupled the single cell interferon response phenotype to the total communication stimulus each cancer received from across the TME via each receptor. The cancer cell data was subset by subclonal cancer genotype (identified in ^10^) and communication scores were square root transformed, scaled and centered (mean=0, sd=1) to improve normality and comparability respectively.

For each cancer subclone of each tumor sample, we identified communications strongly associated with activation of an interferon response, using a lasso penalized likelihood regression model (R package ‘glmnet’). The lasso penalty (α = 1) encourages detection of the communications most strongly activating interferon response, through a shrinkage of the coefficients of all but dominant communications predictors. This variable selection approach minimizes overfitting when considering the role of many communication pathways and enhances the interpretability and predictive accuracy of the model.

Cross-validation (internal 10-fold) was performed to determine the penalty parameter (λ) that minimized the mean cross-validated error. The contribution of each communication to (coefficients) the explained variance in cancer cell interferon phenotype was then assessed. We identified the communication receptors of cancer cells detected to contribute to the interferon phenotype in more than 10% of tumor subclones.

The association of a cancer interferon response with the most communication via the most frequently detected receptor (IL-15RA) was examined using a generalized additive model with a unique smoothing term for resistant and sensitive tumors given each treatment. The IL-15 communication received by cancer cells was also discretized into deciles of signal strength and the distribution of cancer interferon response phenotypes compared to the signal received.

### CD8+ T-Cell isolation and activation

Leukocyte Reduction System (LRS) cones were obtained from two blood donors at City of Hope, Duarte, CA under Institutional Review Board (IRB # 17387) approval. Blood from LRS cones was transferred to K2EDTA blood collection tube (BD Biosciences), and centrifuged for 10 minutes at room temperature and 800 x g. Plasma was removed and buffy coat was collected and diluted to 5mL in 1x PBS without MgCl_2_ (Gibco) + 2% hiFBS. Red blood cells were removed by immunomagnetic depletion using 50μL of EasySep RBC Depletion Reagent (Stemcell Technologies) per mL of sample according to manufacturer instructions. Subsequently, CD8+ T cells were isolated using EasySep Human CD8+ T Cell Isolation Kit (Stemcell Technologies) by immunomagnetic negative selection according to manufacturer instructions.

Isolated CD8+ T cells were centrifuged for 5 minutes at room temperature, 300 x g. T cells were then resuspended to 0.5e6 cell/mL in RPMI-1640 (Gibco) + 10% heat inactivated FBS (hiFBS, Sigma-Aldrich) + 1x antibiotic-antimycotic (Gibco) and stimulated for activation for 3 days supplemented with 20ng/μL IL-2 (Miltenyi Biotec) and CD3/CD28 Dynabeads Human T-Activator (Gibco) in a 6-well tissue culture treated plate (Corning) and maintained in 37°C humidified incubator + 5% CO_2_. Prior to co-culture with cancer cells, CD8+ T cells were collected and CD3/CD28 beads removed using DynaMag-15 magnet (Gibco). Purified activated CD8+ T cells were centrifuged at 300 x g 5min, at room temperature and resuspended to > 2.0e6 cell/mL in fresh RPMI-1640 complete culture media.

### T cell Viability Assay

To assess T cell proliferation in the absence of cancer cells, activated CD8+ T cells were cultured in RPMI-1640 + 10% hiFBS + 1x antibiotic-antimycotic with control (0.1% DMSO), 20ng/μL IL-15 (R&D Systems), 20ng/μL IL-18 (R&D Systems), 5μM ribociclib (Selleck Chemicals), or combinations 20ng/ μM IL-15 + 5μM ribociclib and 20ng/ μM IL-18 + 5μM ribociclib in ULA spheroid plates. T cell growth was monitored using Cytation 5 by brightfield imaging every 12 hours. After 3 days, proliferation was assessed by measuring total ATP using the CellTiterGlo Luminescent Cell Viability Assay (Promega Corporation). We analyzed how T cell viability was impacted by ribociclib and cytokine treatments, individually or in combination. A hierarchical regression model measured the effect of each treatment and combination on log total ATP at 72 hours. A random intercept component was used to account for donor T cell specific baseline ATP levels. Significant treatment effects on T cell viability were determined using two-tailed t-tests.

### Breast cancer cell line culture

Four Estrogen-receptor-positive (ER+), HER2-breast cancer cell lines (CAMA-1, MCF7, T47D, and MDA-MB-134) were lentiviral labeled with Venus fluorescent protein (previously described in ^92^). This approach allowed cancer abundance to be quantified in monoculture or when co-cultured with T cells, which were unlabeled. Cancer cell lines were confirmed negative for mycoplasma contamination using MycoAlert PLUS Mycoplasma detection kit (Lonza) and cell-lines authenticated by STR profiling. All cell lines were cultured and maintained in a 37°C humidified incubator + 5% CO_2_. CAMA-1 and MCF7 cell lines were cultured in DMEM (Gibco) + 10% hiFBS + 1x antibiotic-antimycotic, while MDA-MB-134 and T47D cell lines were cultured in RPMI-1640 + 10% hiFBS + 1x antibiotic-antimycotic.

### Cancer– T cell spheroid co-culture

Cancer cells of each cell line were plated at 1.0e4 cells per well in a total volume of 100μL in 96-well Black/Clear Round Bottom Ultra-Low Attachment (ULA) Surface Spheroid Microplate (Corning) in respective cell line complete culture media. After 24 hours of spheroid formation, 2/3 replicates were treated with 50μL of 0.2e6 CD8+ T cell per mL medium (final of 1.0e4 CD8+ T cells per well) (cancer + CD8+ T cell co-cultures). T cells from each of the two different blood donors were used in half of these replicates (1/3 replicates per donor). The remaining 1/3 replicates were treated with 50μL of culture media (cancer monocultures= no T cell controls). For each mono- and co-culture combination, controlled and replicated experiments (n=3-4) were performed to examine the individual and combined effects of cell cycle treatment (ribociclib) and cytokine treatment (IL-15/IL-18). We prepared 50μL of 4X drug stocks to achieve the following final 1X cytokine/ribociclib treatment combinations in respective cell line complete culture media: control (0.1% DMSO), 20ng/μL IL-15, 20ng/μL IL-18, 5μM ribociclib, or combination 20ng/μL IL-15 + 5μM ribociclib and 20ng/μL IL-18 + 5μM ribociclib. Cancer – T cell co-culture plates were maintained in a 37°C humidified incubator + 5% CO_2_. After 4 days of treatment, a 50% media change was performed with fresh 1X drug stocks using Fluent 780 automated workstation (Tecan).

### Spheroid imaging

Cancer cell abundance was observed throughout treatment by imaging mono-cultures and co-culture spheroids using a Cytation 5 cell imaging multimode reader (BioTek Instruments) under dual Bright Field and YFP (ex 500 / em 542 for Venus florescence) image acquisition using a 2×2 montage image with 5-slice, 50μm z-stack using Gen5 software (BioTek Instruments, version 3.10.06). Spheroids were imaged prior to co-culture addition or ribociclib/IL-15 treatment and then every 12 hours post ribociclib/IL-15 treatment for 3 days and then imaged every 24 hours for a total of 7 days. Raw images were analyzed using Gen5 software including image stitching, Z-projection using focused projection, and spheroid size analysis assessing cancer cell viability as measured by total YFP signal intensity within the calculated spheroid area.

### Spheroid co-culture growth response modelling

For each replicate spheroid, we quantified its speed growth or shrinkage over a 75-hour period following the transient T cell activation phase (*t*_0_= 1,2, 4 and 4 days for CAMA1, T47D, MCF7 and MDAMB134 respectively). For each spheroid, the serial measurements of cancer abundance throughout treatment were scaled relative to the initial cancer abundance at t0. For each cancer spheroid, the relative growth rate (rgr) of each cancer population was determined by the average hourly change in log cancer abundance between the start *t*_0_ and end of treatment *t*_75_, calculated as: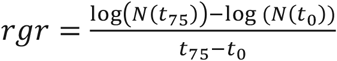 (described in ^93^).

We analyzed how cancer spheroid growth rates (rgr) were impacted by the individual and combined effects of: i) co-culturing with T cells, ii) cell cycle inhibitor treatment with ribociclib and iii) cytokine treatments with IL-15/18, using a hierarchical regression model. The differences in the speed of growth of the four cell lines were accounted for by a cancer lineage specific random intercept component. Similarly, a second random intercept component was added to accounted for donor T cell specific differences in activity across treatments. The full model regression model contained a linear predictor describing cancer rgr based on the effects of: i) CD8+ T cell co-culturing, ii) cell cycle inhibitor treatment and iii) cytokine treatment with either IL-15 or Il-18 and included the three-way interaction terms between these effects. Significant treatment and T cell co-culture effects on cancer growth were determined by comparing the likelihood of the full model, allowing synergies between treatment and co-culturing effects, with nested null models in which: i) treatment and co-culturing effects are additive or ii) treatment or co-culturing have no effect on cancer growth rate. Likelihood ratio tests were used to compare models and identify the factors impacting cancer population growth. The significance of treatment effects, co-culturing effects and synergies were determined using two-tailed t-tests with the Satterthwaite method applied to perform degree of freedom, t-statistic and p-value calculations.

## Author Contributions

J.I.G. contributed study design and coordination, performed mathematical/statistical analyses to evaluated tumor composition, cellular phenotypic heterogeneity and tumor-wide communications related to patient outcomes using scRNAseq data, analyzed in vitro experimental data and wrote manuscript. P.C. performed scRNA and in vitro experiments and wrote manuscript. E.C.M., A.N. conducted bioinformatics pipelines to process scRNAseq data and performed normalization and cell type classification. F.R.A. developed analyses and models and contributed to writing the manuscript. J.T.C. developed bioinformatics pipelines, performed data management and curation and wrote the manuscript. Q.J. K., conceived and coordinated the clinical trial, contributed clinical support and infrastructure and provided clinical data and patient samples as well as contributed to writing the manuscript. A.H.B. designed the research project and analyses, performed scRNA experiments and data analysis, coordinated genomic and mathematical/statistical analyses, and wrote the manuscript.

## Acknowledgements

We thank the anonymous patients from the trial that made this study possible. We thank Anne O’Dea, Priyanka Sharma, Cynthia Ma, Meghna Trivedi, Kevin Kalinsky, Kari B. Wisinski, Ruth O’Regan, Issam Makhoul, Laura M. Spring, Aditya Bardia, Yuan Yuan, Lauren Nye, Onalisa Winblad, Jamie Wagner-Berbel, Kelsey Larson, Christa Balanoff, Gregory Crane, Fang,Fan, Allison Aripoli, Amanda Amin, Richard McKittrick, Marc Hoffmann, Marc Inciardi, Cory Bivona, Mia Hard, Manana Elia, and Mark Redick for conducting the trial and contributed patient samples. We thank Adam Cohen and Brad Nelson for respectively providing clinical and immunological insights. We thank Eleni Farmaki and Vince Grolmusz for providing lentiviral labelled cell lines, Benjamin Copeland for assistance in patient biopsy sample processing and Ben Decato for initial analysis of the raw scRNAseq data. J.G., A.H.B., P.A.C. A.N., F.A. and J.C. were supported by the National Cancer Institute of the National Institutes of Health (NIH) under award number U54CA209978. The content is solely the authors responsibility and does not necessarily represent the official views of the NIH. The High-Throughput Genomics Shared Resource was supported by the NIH Award Number P30CA042014. J.T.C. was supported by a Cancer Prevention Research Institute of Texas Core Facility Support Award (RP170668).

## Code availability

Custom code used in analyses and to produce Figures 1–6 are available on GitHub at https://github.com/U54Bioinformatics/FELINE_project/FELINE_immune_communication.

## Data availability

Raw single cell RNA-seq data are available through GEO under accession code GSE211434 at https://www.ncbi.nlm.nih.gov/geo/query/acc.cgi?acc=GSE211434. The following secure token has been created to allow review of record GSE211434 while it remains in private status: axujqkyupluxbmp. All other data supporting the findings of this study are available from the corresponding author on reasonable request.

