## Supplementary Information for "Cancer cells communicate with macrophages to prevent T cell activation during development of cell cycle therapy resistance"

**Affiliations:**

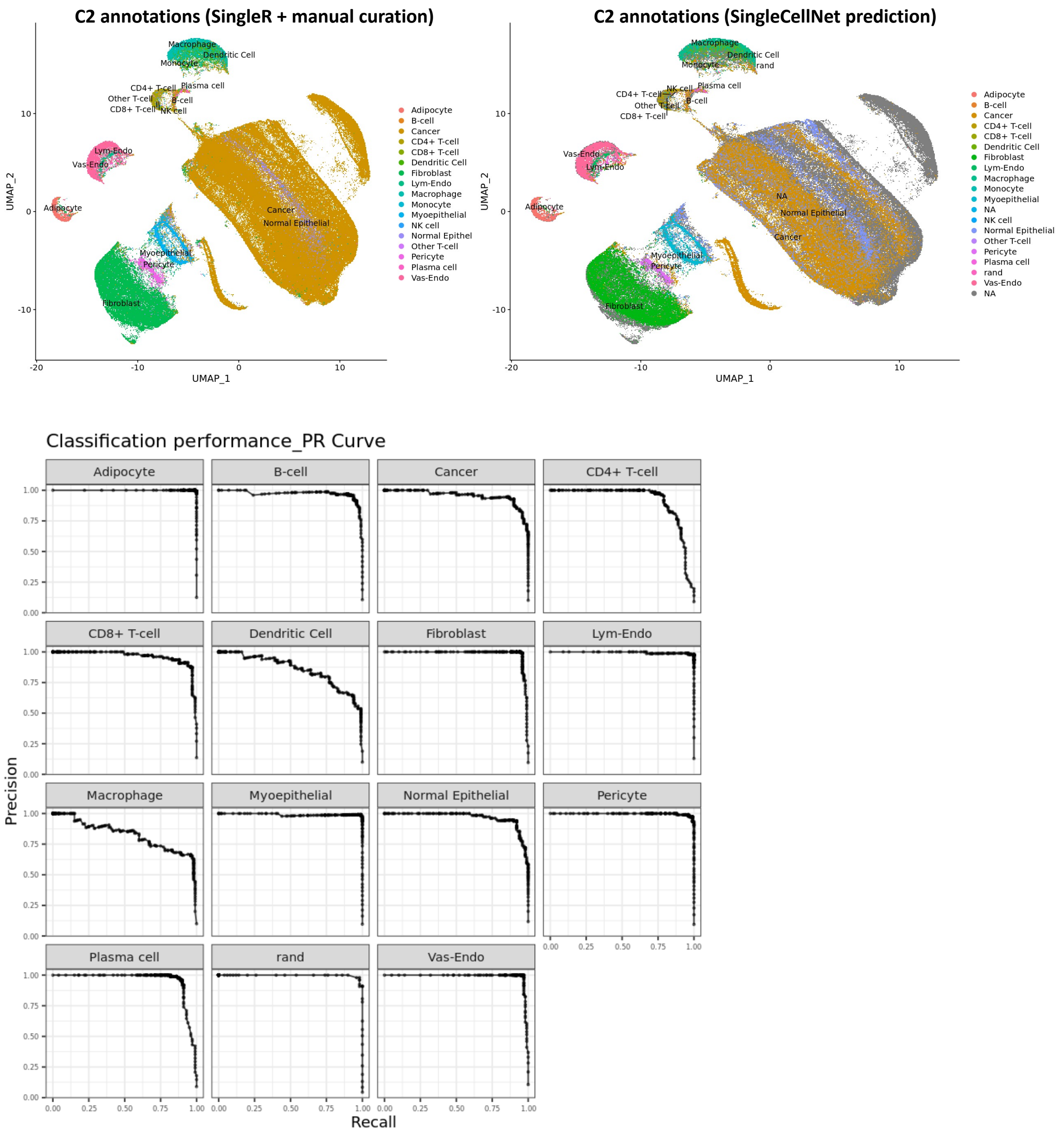


**Figure S1) Validation cohort cell annotations can be predicted using both machine learning of the cell type transcription profiles observed in the discovery cohort or using independent SingleR annotation and manual curation.** This UMAP projection shows high-quality discovery cohort cells grouped by cell type predicted from their gene expression profiles. The UMAP was calculated from the first 10 PC's of the scaled expression of 100 marker genes associated with each cell type. The top left panel shows cells grouped and annotated using SingleR and manual curation while the top right panel shows cells grouped and annotated using predictions of SingleCellNet random forest machine learning classifier. The bottom panel shows the precision recall curves of the random forest classifier in the hold-out validation subset of high-quality discovery cohort cells.

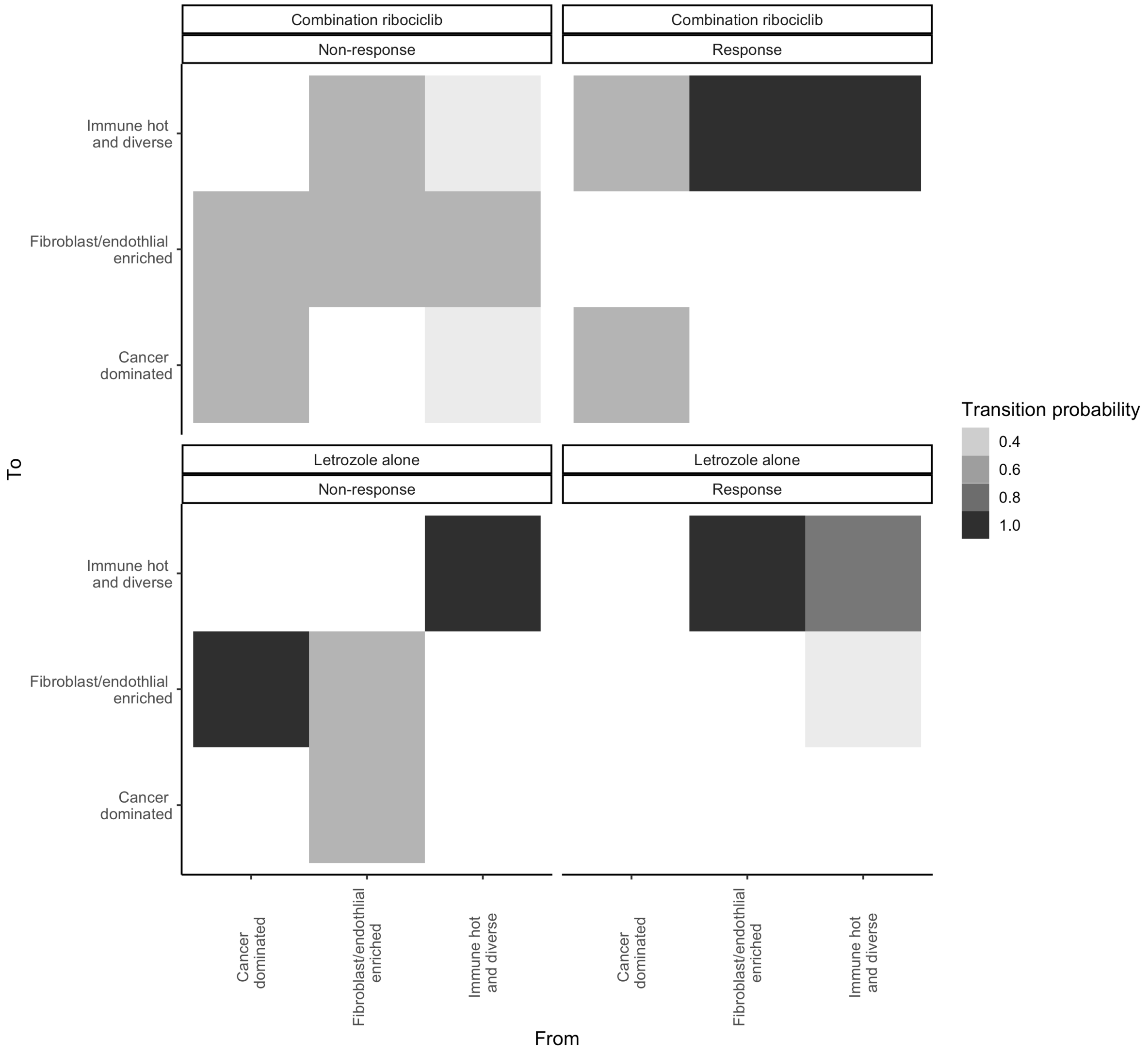


**Figure S2) Ribociclib sensitive tumors get hot by the end of treatment independent of the starting state. However, resistant tumors remain in the cancer dominated or fibroblast/endothelial enriched states.** Heatmap showing the probability of resistant and sensitive tumors receiving combination ribociclib or letrozole alone transitioning between each archetypal tumor composition state. Coloration indicates the proportion of cells transitioning from one state (x axis) to another (y axis).

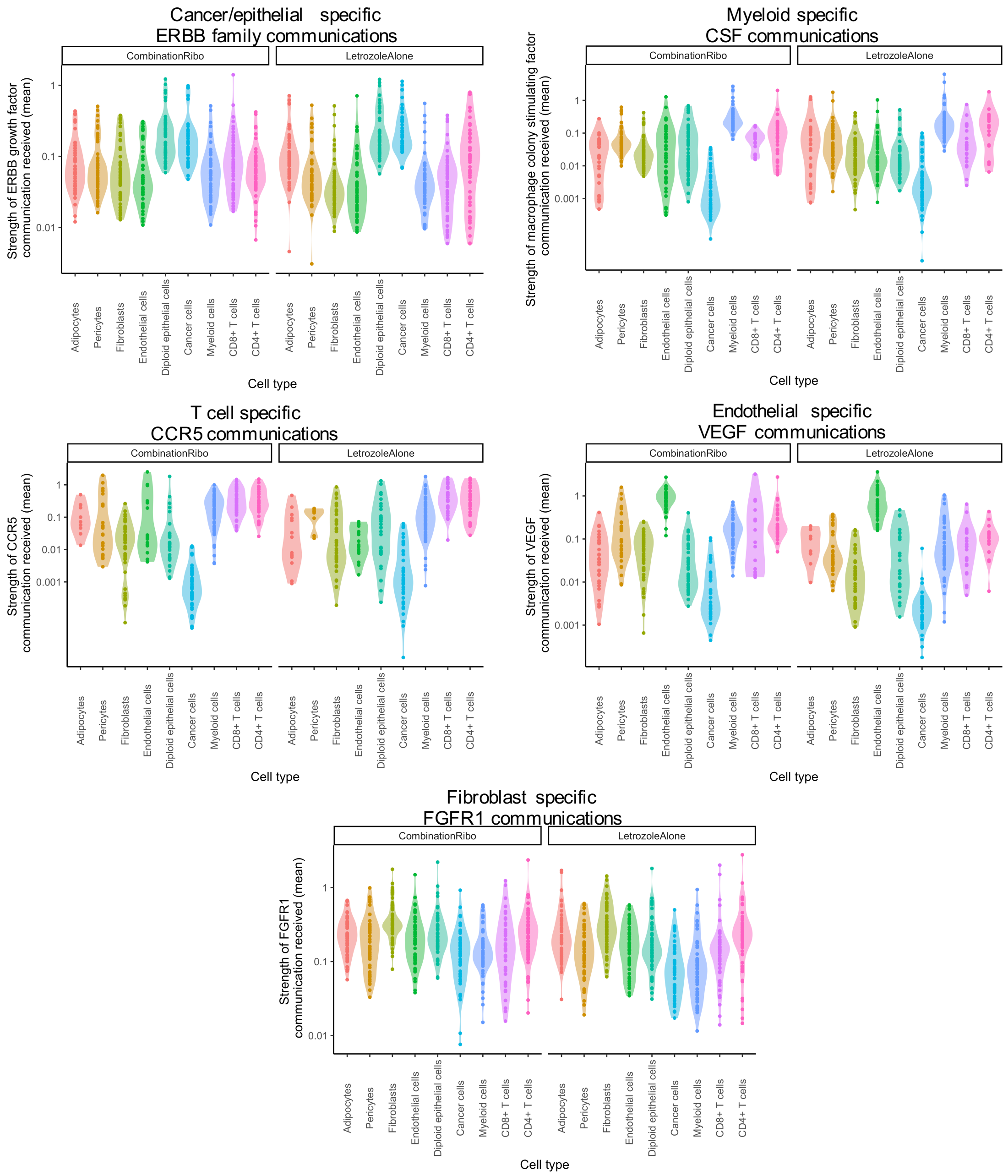


**Figure S3) Validation that TWISTER recovers known communications specifically received by a certain cell type and providing essential signals for activation (e.g VEGF is an angiogenic factor and essential growth factor for vascular endothelial cells).** Violin plots show the log scale strength of communication (y axis) received by cells of each cell type (x axis: color) through 5 cell type specific communications. As expected, epithelial (normal and cancer) cells received higher communications via ERBB family receptors (EGF+ERBB2-4), macrophages received higher communications via macrophage colony-stimulating factor 1 receptor (CSF1R), T cells received higher communications via C-C chemokine receptor type 5 (CCR5), endothelial cells received higher communications via vascular endothelial growth factor (VEGF) and fibroblasts received higher communications via fibroblast growth factor receptor 1 (FGFR1). Each cell type specific communication was found to be significantly activated in the expected cell type (all $p<0.005$).

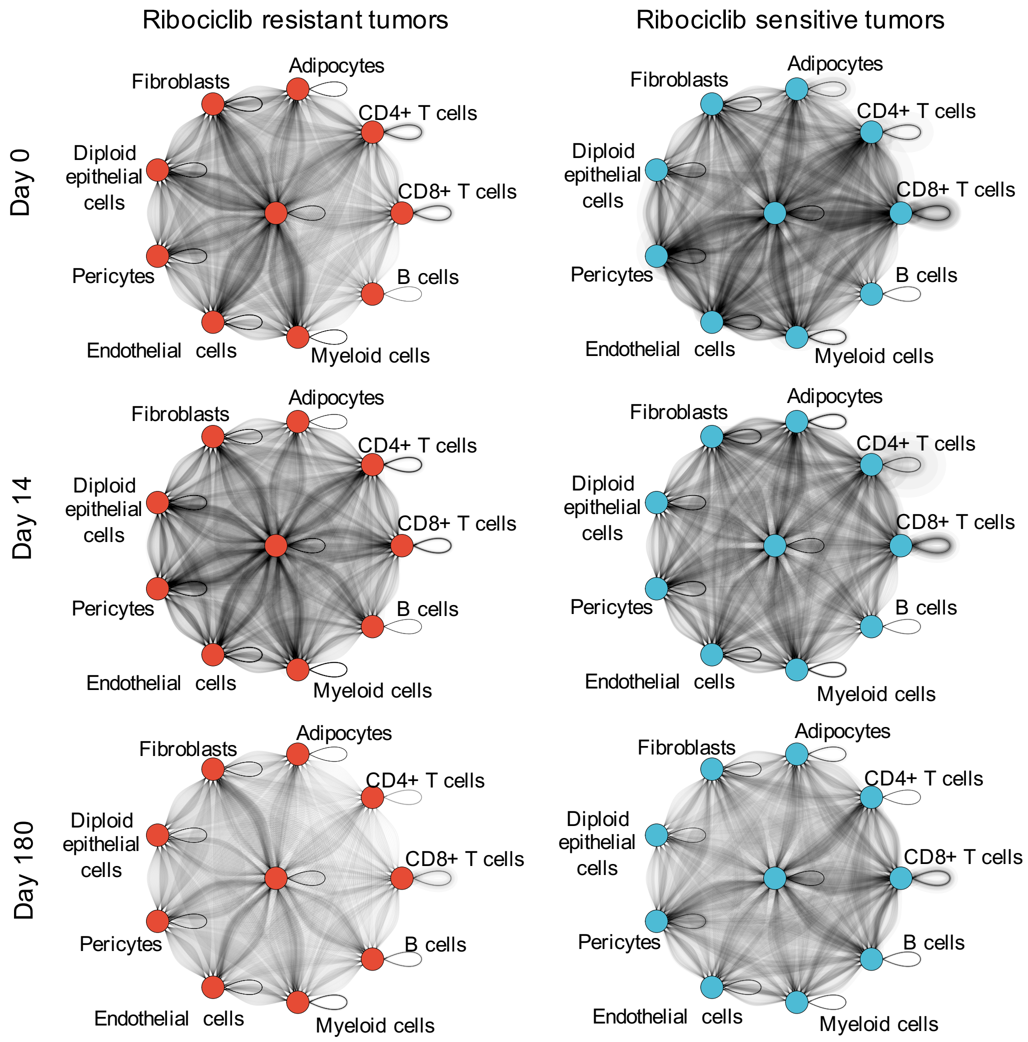


**Figure S4) Global communication differences between ribociclib resistant and sensitive tumors.** TWISTER measurements of tumor wide communication between cell type populations before (top row) during (middle row) and after treatment (bottom row) in ribociclib resistant (left column) and sensitive (right column) tumors. Communication networks show cell types (nodes) and their strength of communication with each other via each LR communication pathway (edges). The darkness of the edge lines is proportional to the strength of communication via that pathway. All edges are directed from one cell type to another, going in a clockwise direction from the sender and to the receiver.

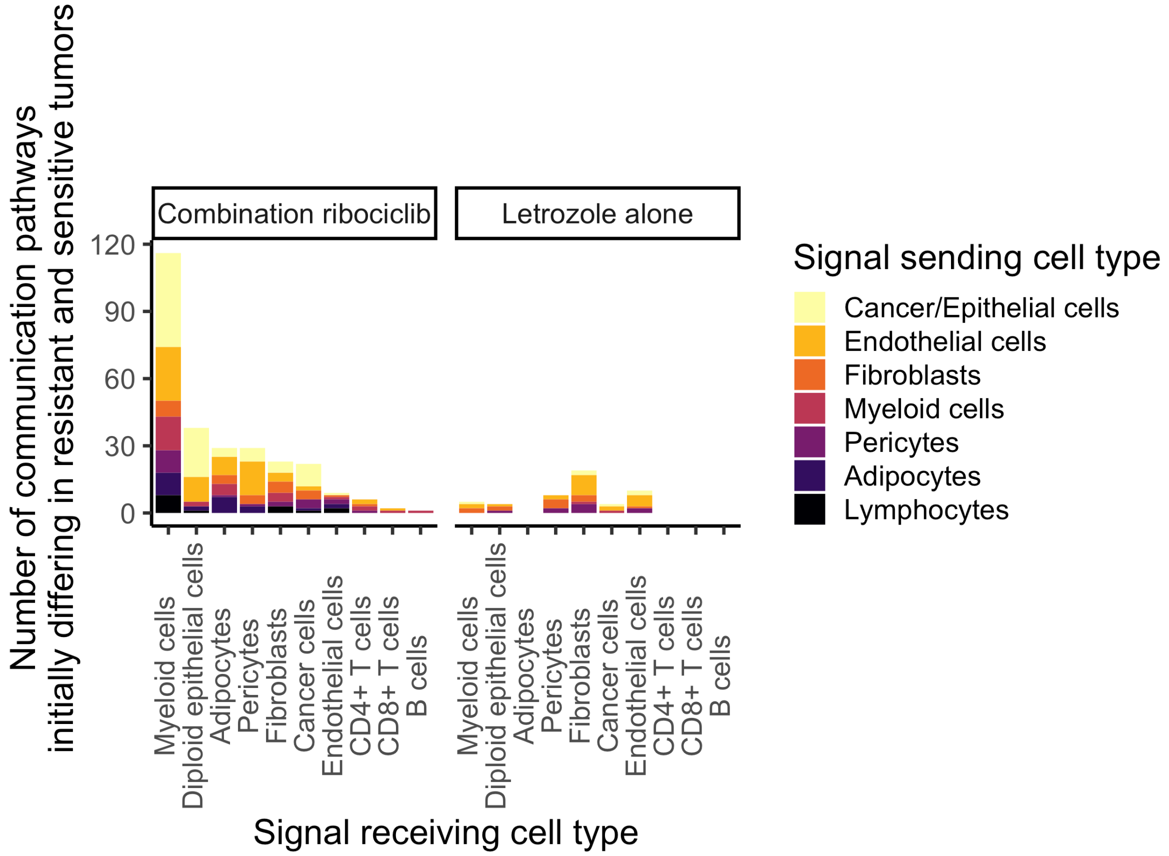


**Figure S5) Frequent communication differences between cell types in tumors resistant and sensitive to treatment.** The y-axis indicates the number of pre-treatment communications that each cell type receives from each other cell type that differ between tumors resistant and sensitive to each treatment (panels). Colors indicate which cell type is sending the signals. The left panel shows the number of pre-treatment dysregulated communications from cancer/epithelial cells to myeloid cells in resistant tumors (all communication pathways detected by ANOVA with adjusted p-values <0.05). The right panel shows the contrasting differences in communication seen between letrozole resistant and sensitive tumors, with fibroblasts being the target of most dysregulated communications.

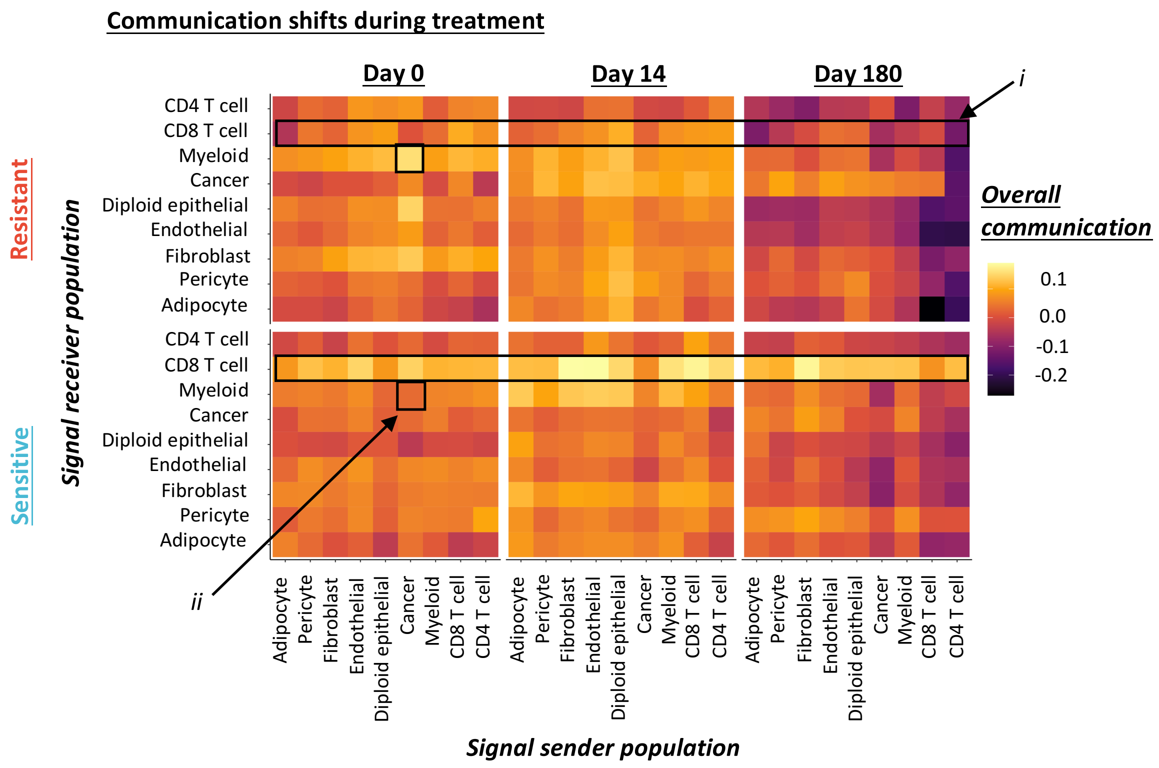


**Figure S6) Summarization of communication measurements reveals changes in the overall communication networks in tumors sensitive and resistant to cell cycle inhibitor therapy (rows) before during and after treatment (columns).** Heatmap shows the pairwise communication between cell types during treatment. Overall communication measures the average signaling between one cell type and another across all L-R communication pathways. Coloration indicates the strength of communication (coloration) from one cell type (x axis) to individual cells of each other cell type (y axis). Boxes indicate: i) the loss of communication with CD8+ T cells during treatment in ribociclib resistant tumors, ii) the stronger communication of cancer cells with myeloid cells prior to treatment in ribociclib resistant tumors.

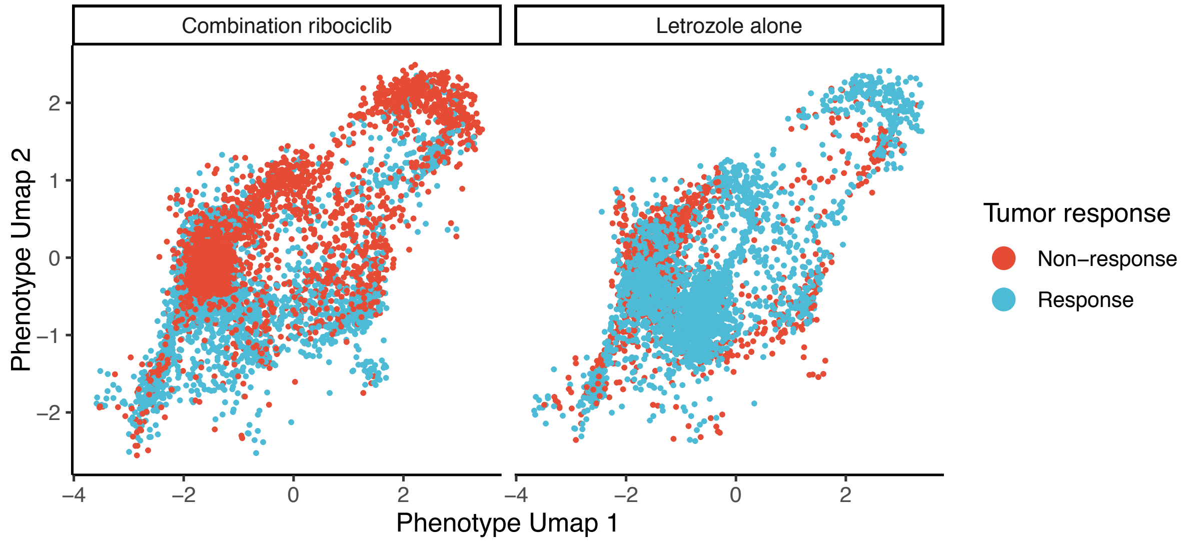


**Figure S7) Verification that the myeloid cells of ribociclib resistant tumors of the independently analyses validation cohort are more differentiated towards a pro-tumor M2 phenotype that those from ribociclib sensitive tumors.** Myeloid cells of the validation cohort were projected into the same umap phenotype landscape as those in the discovery cohort, using the discovery cohort's fitted umap model. This preserves the orientation of cellular phenotype heterogeneity between cohorts, allowing comparison of myeloid cell phenotypic similarity (close cells are phenotypically similar). Higher phenotype Umap 2 scores indicate greater M2 differentiation. Colored points indicate resistant (red) and sensitive (blue) myeloid cells of the validation cohort. As seen in the discovery cohort, the M2 polarization of myeloid cells in resistant tumors was only present under combination ribociclib treatment (left panel) and not letrzole alone (right panel).

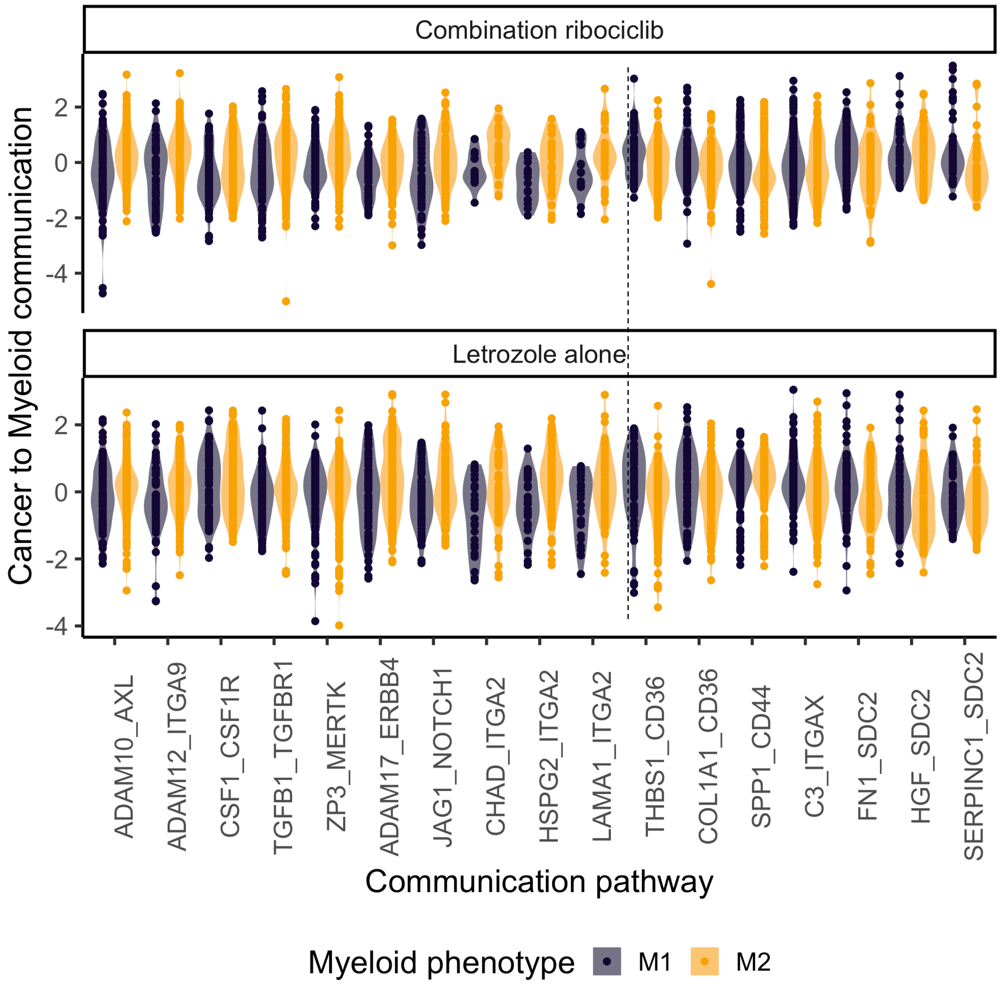


**Figure S8) Established M2 differentiation signals including CSF1-CSF1R, ADAM10-AXL and ZP3-MERTK were sent by and targeted to M2 macrophages.** Cancer communications targeting M2 macrophages were identified by analyzing the average strength of communications from individual cancer cells to macrophages polarized to either an M1 or M2 phenotype (above vs below average differentiation). The LR communications that cancer cells used to differentially communicate with each macrophage phenotype were detected using linear models. Across treatments, cancer cells communicated more strongly with M2 macrophages using communication pathways on the left-hand side of the dashed vertical line and communicated more with M1 macrophages using pathways on the right hand side (all differences in communication significant across treatments with $p<0.005$).

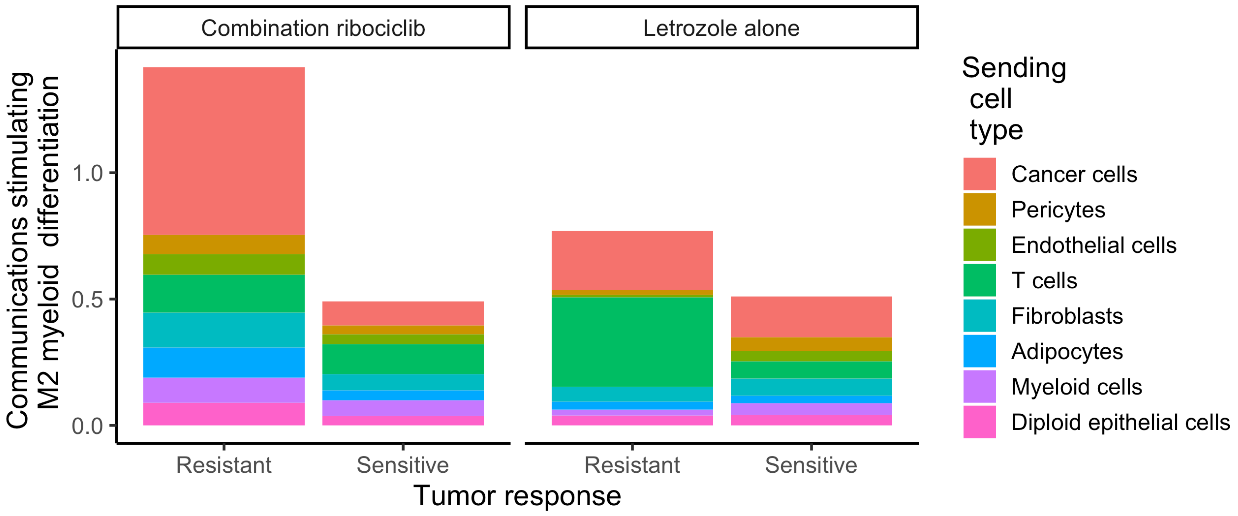


**Figure S9) Cancer cells are the primary contributors of the pre-treatment (Day 0) M2 macrophage differentiation communications promoting the myeloid phenotype shift seen in ribociclib resistant tumors.** For tumors resistant and sensitive to each treatment, we calculated the average pre-treatment contribution of each cancer and non-cancer cell type to the M2 differentiation communications received by macrophages (y axis) in resistant versus sensitive tumors (x axis), totaling the signals across scaled LR pathways. Cancer cells were found to be the dominant contributors to M2 differentiation communications in ribociclib resistant tumors (left panel). The tumor-wide strength of communication (total bar height) was stronger in resistant tumors, with the cancer cell contribution (left side cancer section) surpassing that of all cell types in the sensitive tumors (total height of right bar. This indicates that the M2 macrophage differentiation in ribociclib resistant tumors was driven primarily by corrupting cancer communications that evolved prior to treatment.

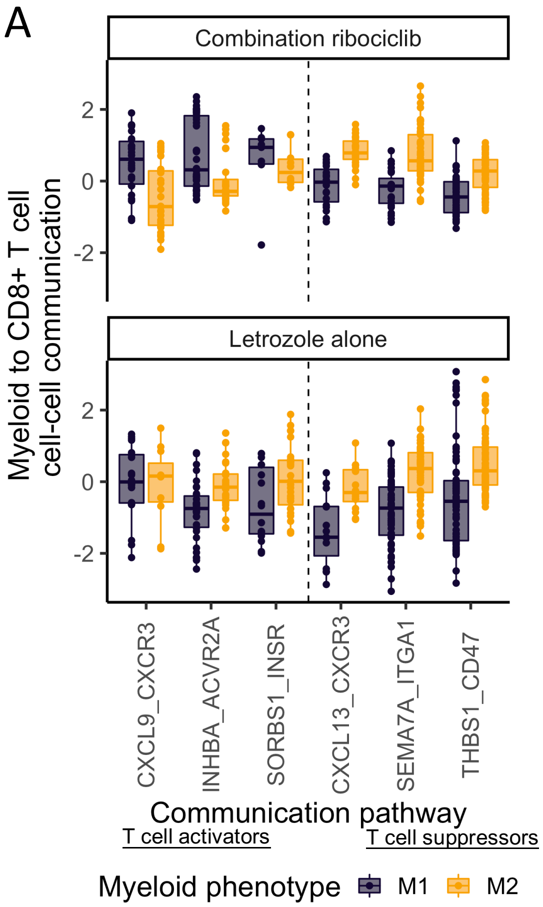


**Figure S10) Verification that M1 macrophages send primarily immune activating signals whereas M2 macrophages send more immune suppressing signals to CD8+ T cells.** Boxplots show the differential communication of M1 and M2 myeloid cells with CD8+ T cells under each treatment (rows). The M1 and M2 macrophage specific communications with CD8+ T cells were identified by comparing individual cell-cell communication, via different LR pathways across tumors. Hierarchical random effects models detected communications differentially sent by individual M1 or M2 macrophages to CD8 T cells, while controlling for the activation state of the signal receiving T cell. Multiple comparisons were accounted for using FDR correction. As expected, the M1 macrophages sent stronger immune-activating signals to T cells (left of vertical dashed line) whereas the M2 cells sent immune-suppressing signals (right of vertical dashed line). Notably, M1 macrophages amplified immune-activating signals much more in ribociclib treated tumors compared those given letrozole alone. For example, inhibin (INHBA) communications inhibit the development of a regulatory T cell (Treg) phenotype. This T cell activating communication was more highly activated in tumors receiving ribociclib compared to letrozole alone. Similarly, M1 macrophages amplified inflammatory communications via C-X-C Motif Chemokine Receptor 3 (CXCR3) in ribociclib tumors more than in tumors given letrozole. These results support the central role of macrophages in activating or suppress an antitumor immune response and driving divergent immune compositions and treatment response of resistant and sensitive tumors.

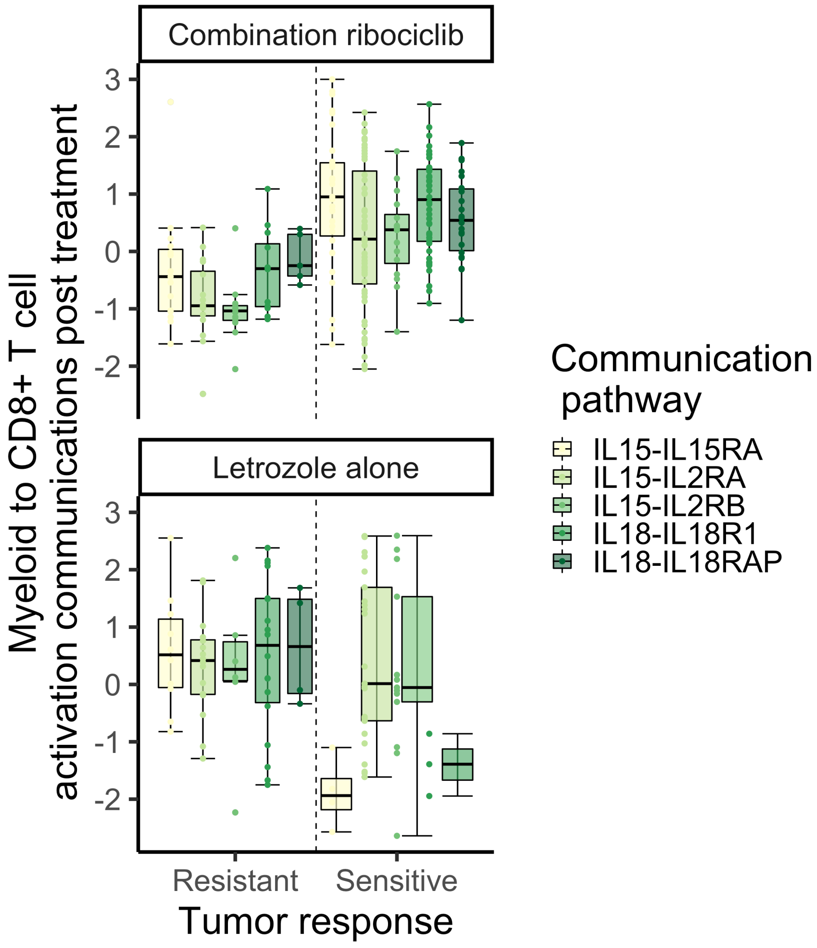


**Figure S11) Verification in the independently profiled validation cohort that post treatment (Day 180) myeloid cells of ribociclib resistant tumors send fewer immune activating communications to CD8+ T cells via IL-15/18.** Box and whisker plots show the weaker CD8+ T cell activating communications received from the myeloid population in combination ribociclib resistant tumors compared to sensitive tumors (top panel right versus left of dashed line) at the end of treatment (Day 180). Colored boxes indicate the CD8+ T cell activating interleukin communications from myeloid cells in tumors of the validation cohort. Communication pathways are the same as those reduced in ribociclib resistant tumors post treatment in the discovery cohort. The IL18-I18RAP communication could not be evaluated under letrozole alone treatment due to a high dropout rate of the IL18RAP receptor.

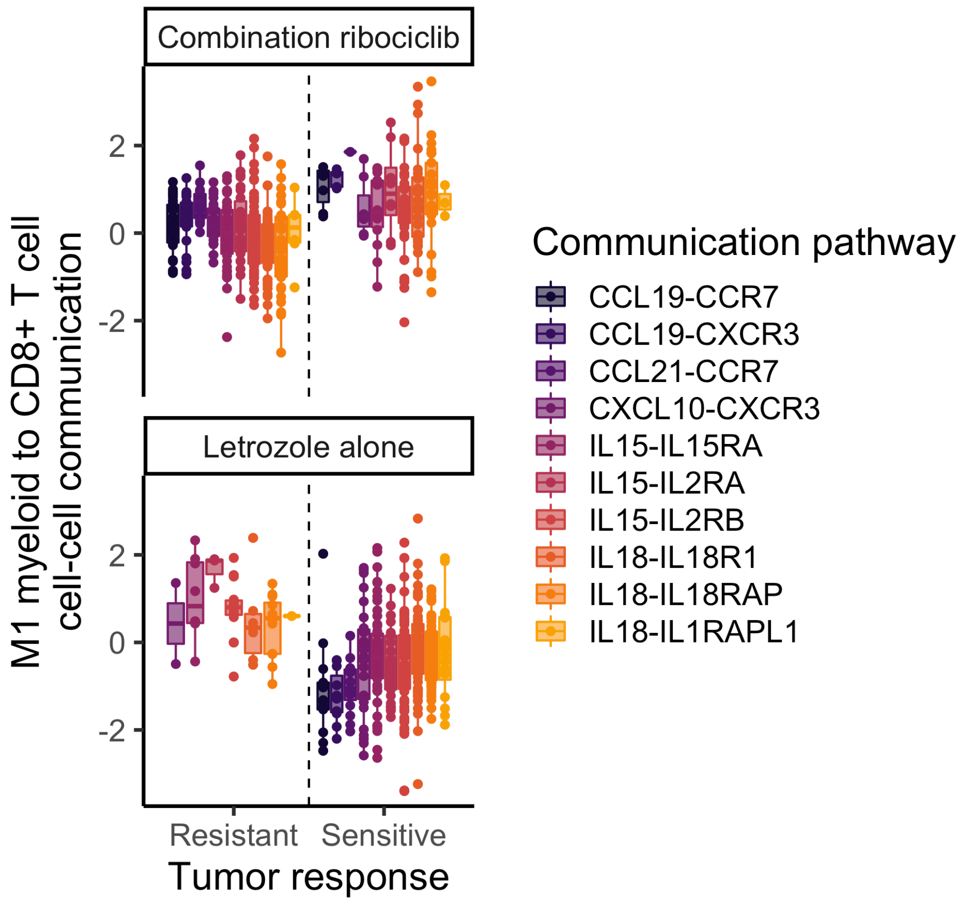


**Figure S12) Suppression of immune activating signals from individual M1 macrophages to CD8+ T cells in ribociclib resistant versus sensitive tumors.** To measure the ability of individual macrophages to activate an immune response, we next contrasted the M1 macrophages cell-cell communications with cytotoxic CD8+ T cells between resistant and sensitive tumors (points=per M1 cell communication with individual CD8+ T cells). Significant differences in cell-cell communication between resistant and sensitive tumors detected using a hierarchical random effects model. In ribociclib resistant tumors (top panel), individual M1 macrophages sent substantially weaker immune-activating signals to CD8+ T cells compared to those of sensitive tumors, including suppression: CCL19/21 stimulation of T cell expansion (df=39, t=3.68, $p<0.001$), CXCL10 promotion of effector T cell differentiation (df=18, t=2.28, $p<0.05$), interleukin 15 (IL-15) stimulation of proliferation and survival (df=111, t=83.3, $p<0.005$) and interleukin 18 (IL-18) activation of an interferon response (df=60, t=4.71, $p<0.0001$). The suppression of M1 immune activating signals was not seen in letrozole resistant tumors (bottom row). These findings reveal how cancer induced polarization of macrophages can suppress an effective immune response by reducing the T cell activating communications sent by individual M1 macrophages.

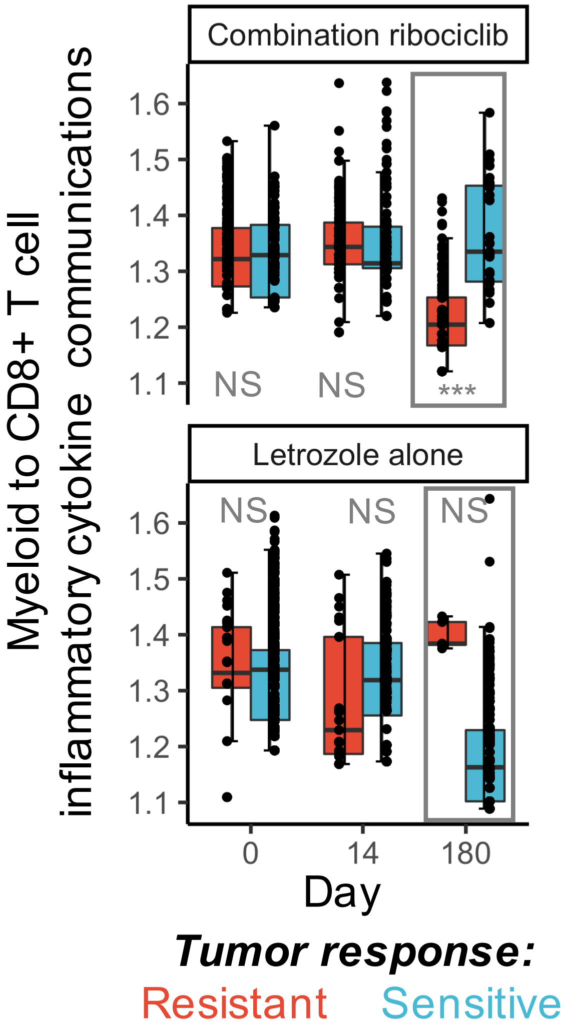


**Figure S13) Suppression of inflammatory cytokine communications from the myeloid population to CD8+ T cells by end of treatment in ribociclib resistant but not sensitive tumors.** Immune activating inflammatory cytokine communication pathways were identified by their receptors using the gene-ontology database. The overall immune activating myeloid to CD8+ T cells communication was measured by totaling the inflammatory cytokine communications from across the diverse myeloid population. We then analyzed how this immune activation communication diverged during treatment in resistant and sensitive tumors. This analysis revealed that the M2 polarized macrophage populations of resistant tumors provided fewer immune activating signals to CD8+ T cells throughout ribociclib treatment (df=185, t=4.48, $p<0.0001$). Conversely, myeloid signaling to CD8+ T cells was maintained in ribociclib sensitive tumors (stronger communication vs resistant tumors: df=778, t=6.79, $p<0.0001$) and the suppression of immune activating CD8+ T cell communications was not seen in the letrozole resistant tumors (df=113, t=-1.83, $p=0.07$).

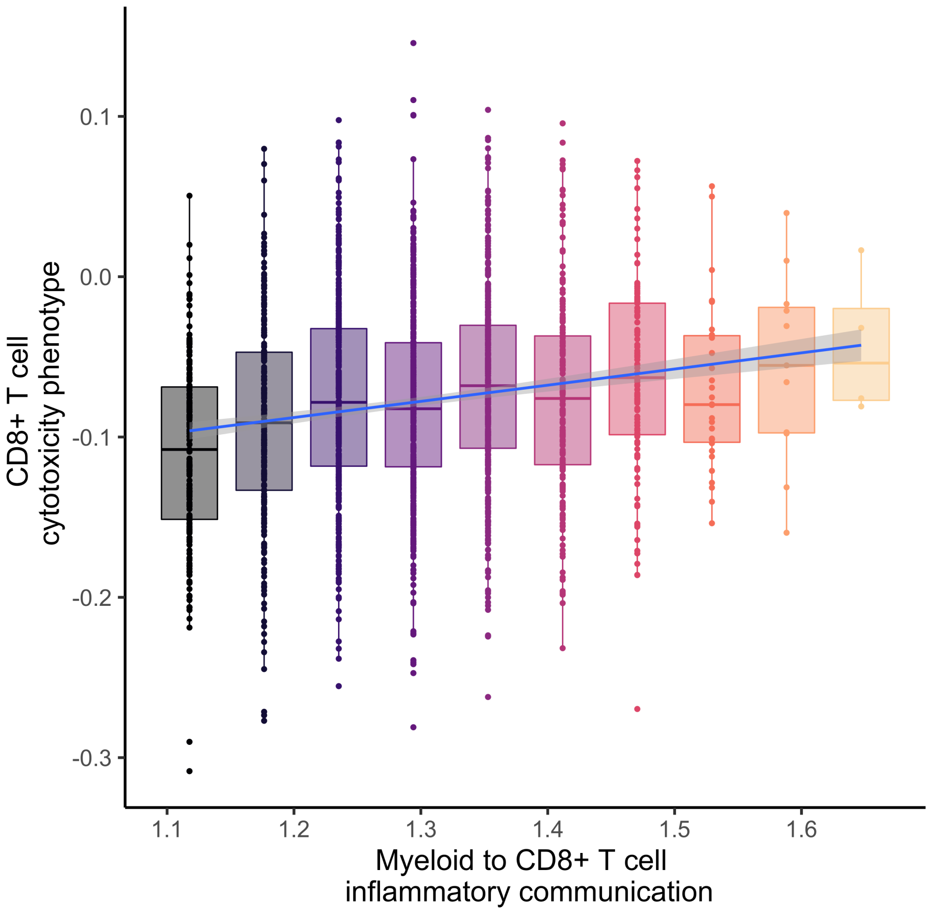


**Figure S14) Stronger myeloid communication of inflammatory cytokine signals supported CD8+ T cell activation to an effector phenotype.** Scatterplot shows the strongly significant association between the inflammatory cytokine communications from macrophages to CD8+ T cells and the differentiation of that T cell towards a cytotoxic phenotype. The T cell differentiation was measured using a CD8 T cell specific ssGSEA pathway contrasting gene expression of naive and cancer killing effector cells (GSE22886 Naive CD8 T cell vs NK cell up). For each CD8+ T cells (points) differentiation state was linked to the inflammatory cytokine communication it received from across the myeloid population. Across both ribociclib and letrozole alone treatments, greater macrophage to T cell inflammatory cytokine communication was associated with activation of an effector T cell phenotype (df=786, t=4.98, $p<0.0001$).

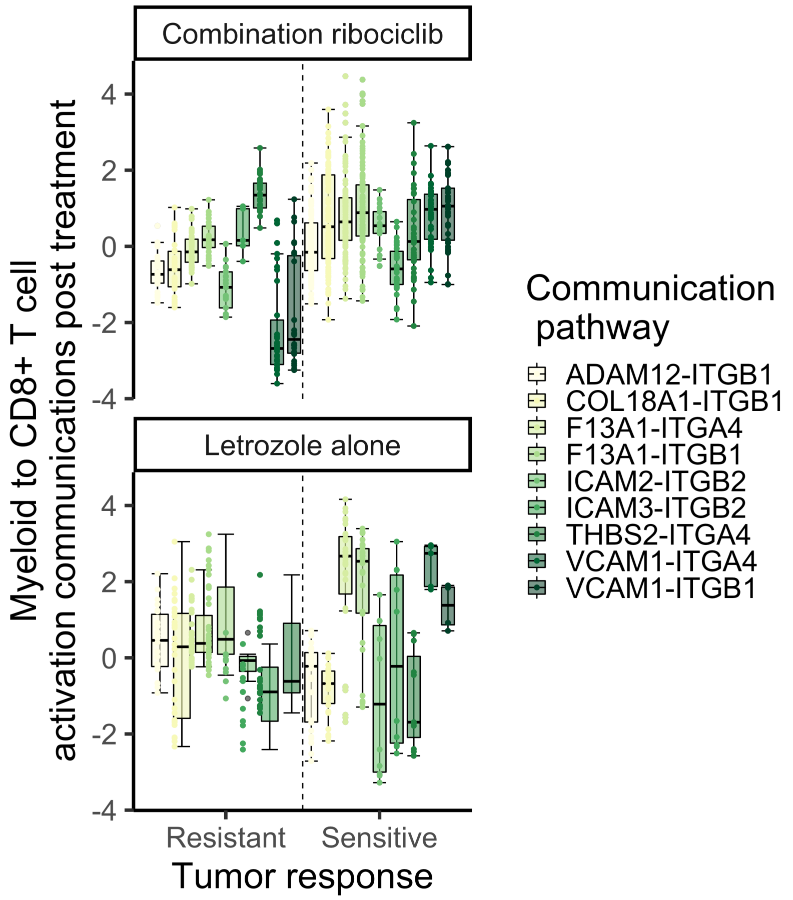


**Figure S15) Verification in the independently profiled validation cohort that post treatment (Day 180) myeloid cells of ribociclib resistant tumors send fewer immune recruiting communications to CD8+ T cells via IL-15/18.** Box and whisker plots show the weaker CD8+ T cell recruitment communications received from the myeloid population in combination ribociclib resistant tumors compared to sensitive tumors (top panel right versus left of dashed line) at the end of treatment (Day 180). Colored boxes indicate the CD8+ T cell recruiting integrin communications from myeloid cells in tumors of the validation cohort. Communication pathways are the same as those reduced in ribociclib resistant tumors post treatment in the discovery cohort.

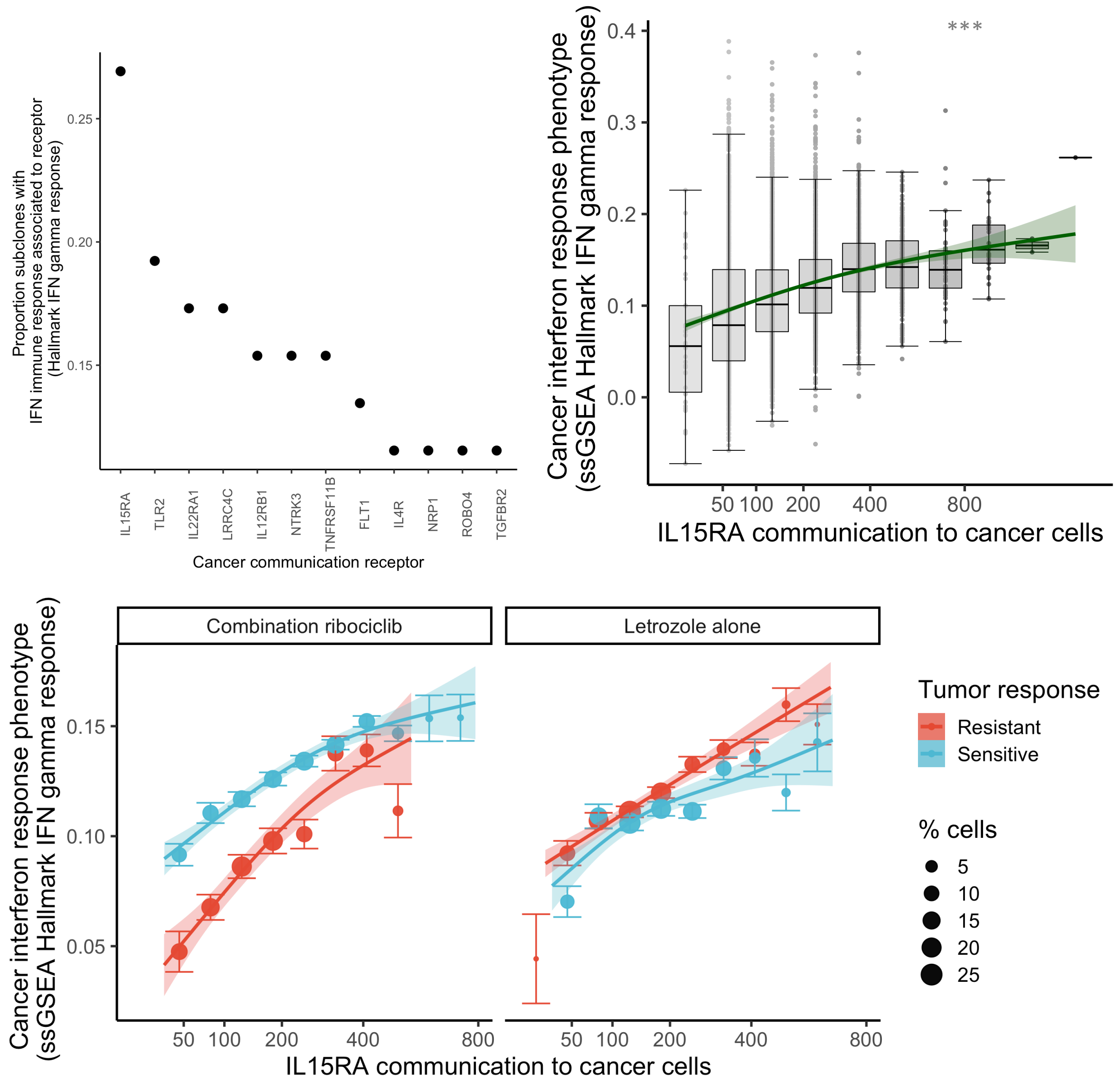


**Figure S16) upp :Lasso Hallmark IFNGamma Cancer predictors) Cancer activation of an interferon gamma immune response post treatment linked to receipt of tumor wide IL15 communications.** We assessed the post treatment (day 180) communications most strongly associated with an immune response phenotype in cancer subclones. We expected that in immune hot TME's, the high levels of immune activating signals, T cell recruitment and activation should induce a phenotypic response of cancer cells. Cytokine signals should be transduced via Jak-STAT, to the cancer nuclei, driving activation of interferon regulatory factors (IRFs) and induction of IFN-stimulated genes (e.g Interferon gamma-induced proteins) and increase the production of antigen presenting major histocompatibility complex molecules (MHC I) allowing recognition and killing of cancer cells (PMID: 22390970; PMID: 28791024). The cancer cell activation of this interferon gamma response pathway was measured using the hallmark interferon gamma response ssGSEA scores to contrast pathways activation across cancer cells. For each subclonal cancer population, we identified communication pathways strongly associated with interferon gamma response phenotype activation, using Lasso regression (see methods).  Induction of a cancer interferon gamma response was most frequently associated with strong Interleukin 15 Receptor Subunit Alpha (IL15RA) communications from across the TME (28% of tumor subclones) (top left panel). Other detected communications impacting the cancer interferon response bound to cytokine receptors such as: Toll-like receptor 2 (TLR2: 19% subclones), Interleukin 22 Receptor Subunit Alpha 1 (IL22RA1: 17% subclones), Interleukin-12 Receptor Subunit Beta-1 (IL12RB1: 15% subclones). The strongly significant association of IL15RA communication and cancer interferon gamma response was consistent across tumors resistant and sensitive to both treatments (top right panel) ($df=788$, $t=-2.90$, $p<0.005$). However, cancer cells from ribociclib resistant tumors received weaker IL15RA communications and these cancer cells exhibited the weakest interferon gamma response (bottom panel) ($df=788$, $t=-10.05$, $p<0.0001$). This lack of immune detection was not observed in letrozole resistant tumors. The result was confirmed in the validation cohort, with resistant cancer cells having lower interferon gamma response than those of sensitive tumors under ribociclib but not letrozole treatment ($df=26211$, t-28.06, $p<0.0001$).

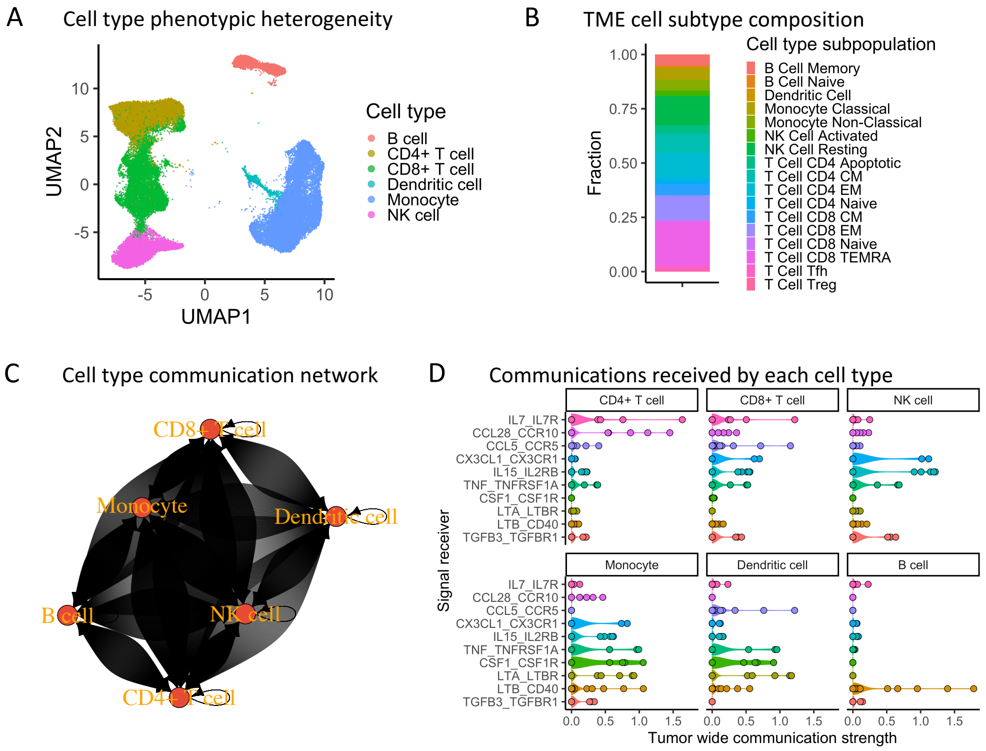


**Figure S17) Recovery of known cell type specific communications between immune populations in the peripheral blood using the TWISTER approach.** Published scRNAseq data from a study exploring the phenotypic evolution of circulating immune cells of gastrointestinal cancer patients during immunotherapy (GEO accession no. GSE130157) was used to validate that known cell interactions can be recovered. A) The transcriptomic profile of the 70781 cells sampled in the study was used to characterize the phenotype heterogeneity of immune cell types, using umap. The phenotypic similarity (little overlap of cell type clusters) within cells annotated as T cells, B cells, monocytes and dendritic cell shows the distinct phenotype of each cell type. B) Using the pre-treatment sample from patient HJD33E we calculated the relative abundance of cell types and the subpopulations of phenotypically distinct cells within each cell type (e.g. naïve, central memory (CM), effector memory (EM) and effector memory cells re-expressing CD45RA (TEMRA) T cells). Distinct cell subtype phenotypes were identified by the authors using differential gene expression analyses. C) TWISTER was used to reconstruct the communication between cell type subpopulations. The directed network graph shows the strength of different ligand-receptor communications sent by a cell type population and received by an individual of another cell type. The darkness of lines indicates the strength of communication (darker=stronger communication via a specific LR pathway). Arrows are directed from one cell type to another in a clockwise direction. Across communication pathways, we observed strong reciprocal communication of monocytes and CD4+ T cells, CD8+ T cells and NK cells. Weaker communication was seen between B cells and Dendritic cells. D) Specific LR communications (y axis) known to be particularly received by certain cell types (panels) were extracted. TWISTER inferred the communication strength received by immune cells of each type from each cell types population (points). Violin plots show the distribution of communication strengths received from across cell types. As expected, T cells received greater IL7 signals, highly activated effector NK cells received greater signals via CX3CR1, monocytes received greater CSF1 and TNF signals and B cells received greater signals via CD40.

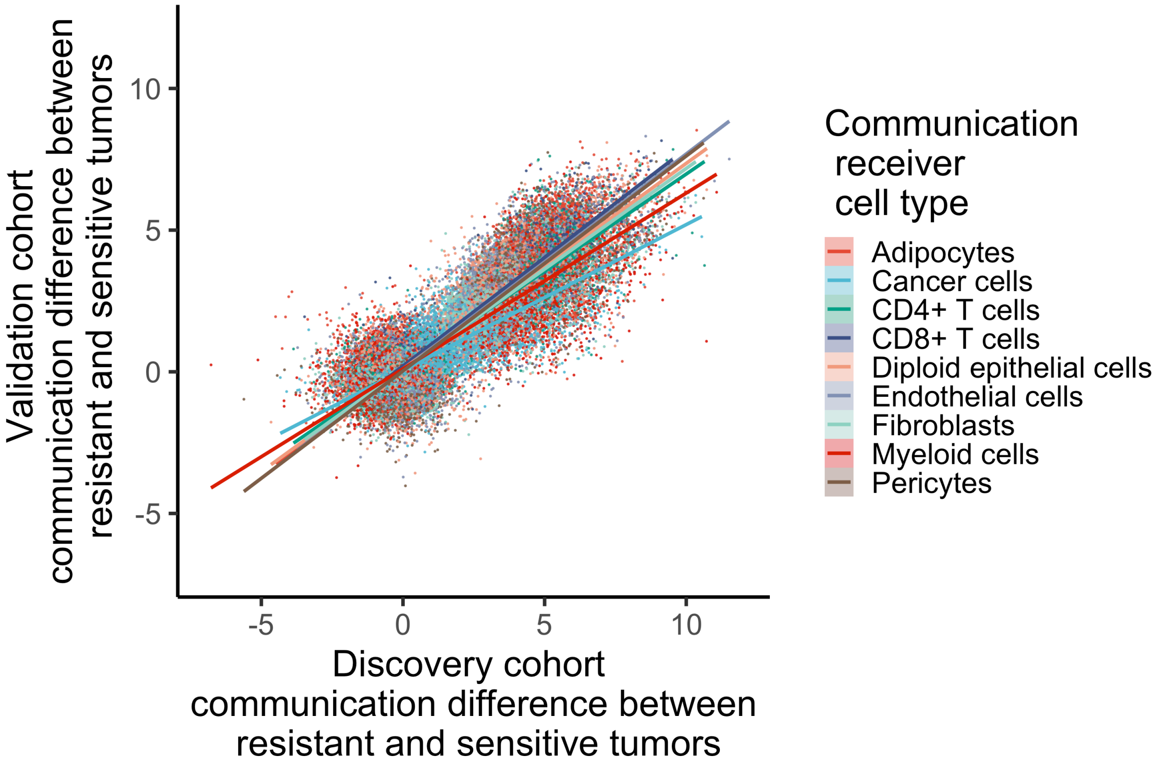


**Figure S18) Consistent communication measurements between the independently profiled discovery and validation cohorts)** Scatterplot comparison of communication measurements between discovery (x axis) and validation (y axis) cohorts. Each point represents an estimate of the strength of communication from one cell type to a focal receiving cell type (color) via a specific LR communication pathway. Regression lines are plotted showing the consistency of the strength of communication measured in the discovery and validation cohort. A strong and significant correlation of 0.82 was found across cell types (df=106734, t= 679.01, p <0.00001)
